# Connecting conformational stiffness of the protein with energy landscape by a single experiment

**DOI:** 10.1101/2020.06.09.142257

**Authors:** Soham Chakraborty, Deep Chaudhuri, Dyuti Chaudhuri, Vihan Singh, Souradeep Banerjee, Shubhasis Haldar

## Abstract

Proteins are versatile biopolymers whose functions are determined by their structures. Understanding the structural dynamicity, with respect to energy landscape, is essential to describe their biological functions. The ability to study the dynamical properties of a single protein molecule is thus crucial, but ensuring that multiple physical properties can be simultaneously extracted within a single experiment on the exact same protein molecule in real-time has hitherto been infeasible.

Here, we present magnetic tweezers technology that surmounts this limitation, providing real-time dynamic information about changes in several molecular properties (*Δ*G^0^, conformation, and mean first passage time of unfolding and refolding) from a single experiment, by *remeasuring the very same protein molecule* in varying chemical condition. We illustrate the versatility of the method by studying electrolyte-dependent conformational flexibility and the energy landscape of substrate protein L under force. Changing salt concentrations reshapes the energy landscape by two specific ways: it speeds-up refolding kinetics while slowing down unfolding kinetics. From the same trajectory, we calculate the stiffness of the protein polymer, a quantity that varies with salt concentration. The data is described within the framework of a modified ‘electrolyte FJC model’ that we propose and study here. The observed correlation between *Δ*G^0^, kinetics and polymer elasticity connect protein chain physics and the energy landscape, while the experimental methodology we describe of getting energy landscape from a single experiment could have wide-ranging applications.

## Introduction

Biomolecules are dynamic in nature and exhibit conformational changes as they interact and transmit downstream signals in the cells^1–5^. Biological processes such as, DNA replication, RNA recognition and folding engage non-interacting regions of the molecules which promote structural flexibility to these molecules^6–8^. Thermodynamics, kinetics and conformational flexibility are the most fundamental molecular properties to determine the nature of these processes. Proteins are also involved in diverse biological processes, ranging from transcription to signal transduction and thus, dynamic changes in these properties are of immense importance to understand their intrinsic structure-function dynamics^1^. The nature of protein interactions, with DNA, ligand, nanoparticles or drugs, are solely dependent on the change in thermodynamic or kinetic parameters^9–11^. To date, these changes are measured either by averaging out a large ensemble of molecules (in bulk assays) or comparing two independent molecules in single molecule technologies. Interestingly, these dynamic changes never been monitored simultaneously, on the exact same protein molecule.

Under single molecule resolution, the effect of ionic strength on elastic behaviour of DNA, RNA polysaccharides and unstructured PEVK domain have been widely studied under high and low force regimes^12–20^. Thermodynamic stability of these biopolymers is largely dependent on the effect of solvent counter-ions that screen their backbone charges. The mechanics and conformational dynamics of DNA, RNA hairpin, DNA-protein interaction, and ion-dependent structural changes in DNA have been monitored under a wide range of tension, both by theoretical tools and by integrating single molecule technologies and microfluidics^21–28^. The advantage of using nucleic acids and polysaccharides as intricate polymers are their higher persistence length and uniformly charged backbone, which aid to detect the conformational stiffness easily in presence of changing salt conditions. Proteins are one of the important biomolecules in the cell and function vastly under varied ionic strength and physiological force. The effect of electrostatic interaction and osmolytes on the mechanical stability of protein GB1 domain have been characterized by force spectroscopy techniques.^29–31^ Similarly, studies have also explored the contribution of different Hofmeister salts to the mechanical stability of proteins under force.^32,33^ However, how varying ionic strength influences structural properties of small globular protein under mechanical constrains has not been reported. This limitation encouraged our present work to study and directly quantify the microscopic chain physics of small globular protein under varying ionic conditions.

Intracellular ion homeostasis plays an indispensable role in maintaining protein conformation by constantly changing ionic environment and thus, directly influence folding^34–36^. Cytoplasmic compartments such as, endoplasmic reticulum, Golgi apparatus or lysosome, involved in secretory pathway have a pH gradient due to the variation in ion compositions which not only affect the folding of the secreting proteins but also of their resident proteins^37^. Additionally, the cytoplasmic ions interact with the proteins and accelerate the folding and stabilize the folded structure by screening dispersed charges in their backbone^38–40^. Charge-dense polymers such as, actin is well known to function under force and is modulated by salts through both nonspecific electrostatic effect and specific binding to their discrete sites^41–43^. Their fundamental properties such as, bending stiffness and interactions with other proteins are also ionic strength dependent^44–50^. However, how the elasticity of these proteins changes as a function of force and electrolyte concentrations is still not known. Though, experimental and theoretical studies suggest that solvent quality affects the compaction of protein polymer, its role in further structure formation to promote folding is highly debated^51^. Additionally, a proper model is lacking, which combines both kinetics and thermodynamics to polymer elasticity in a single frame.

Here, we not only measure several physical properties (*Δ*G, conformation, and mean first passage time of unfolding and refolding) of a single protein but also simultaneously quantify their electrolyte-dependent changes by re-measuring the exact same protein molecule under force. Magnetic tweezers technology allows us to measure the dynamic structural changes of the very same protein molecule in varying chemical condition from a single experiment. In this work, we used ammonium sulfate as a model salt to systematically investigate the effect of ionic strength on the folding landscape and polymer elasticity of protein L B1 domain. Ammonium sulfate, as a protein stabilizer in the Hofmeister series, can effectively collapse the unfolded polypeptide, resulting in more rapid folding^52,53^. Our results show that salts reshape the energy landscape towards the folded state by increasing the height of the free energy barrier with respect to the folded state, keeping the distance to transition state constant. Additionally, it has been predicted that ammonium sulfate as a poor solvent, could facilitate the hydrophobic collapse in the globular protein by modulating their hydrophobicity which bring the non-polar residues closer and leads to a rapid collapse transition^54^. The key finding of our work has been reconciled with a different salt (Sodium Chloride) and protein-talin, which has been well characterized to function under physiological force.^55–58^ Here, we report an ion-dependent protein collapse under physiological force range and propose an electrolyte freely jointed chain (Ec-FJC) model to explain the electrolyte-dependent conformational flexibility in protein molecule. Finally, we successfully combine these results with our kinetics and thermodynamic data to illustrate a comprehensive picture of protein chain elasticity which will open a new gateway for studying folding. Interestingly the data, we observe from our single experiment, are in strong agreement with the multi-experiment observation, claiming their place as a multiplexed biophysical technology to study a wide range of questions in biology.

## Results

### Magnetic tweezers technology to study protein dynamics

Magnetic Tweezers have the advantage of constant monitoring the folding dynamics of a single protein, while introducing external stimuli (proteins/DNA/drugs/nanoparticles etc.) in a single experiment (Fig. 1A). Additionally, it provides us a way to monitor the mechanical unfolding and folding transitions of single protein by applying a *force-clamp technology*. The experiments were performed on polyprotein construct of protein L B1 domain, which was immobilized on glass surfaces using HaloTag-based covalent chemistry and anchored to paramagnetic beads by biotin-streptavidin chemistry. A pair of permanent magnets, placed above the flow chamber, can apply a passive force-clamp upto 120 pN with sub-pN resolution by generating a magnetic field. The pulling force is calculated as an inverse function of distance between magnet and the paramagnetic beads attached to protein and is defined by the magnet law, described by Popa et al (Fig. 1B)^59^. Fig. 1B demonstrates a representative force-clamp trajectory performed by changing magnet position from 4 mm to 1.4 mm, which results in an increase in force pulse from 4.3 pN to 45 pN^59–61^. The applied force leads to eight successive unfolding events observed as increase in step-size (defined as the difference between unfolded state and folded state of a domain).

**Figure 1:**
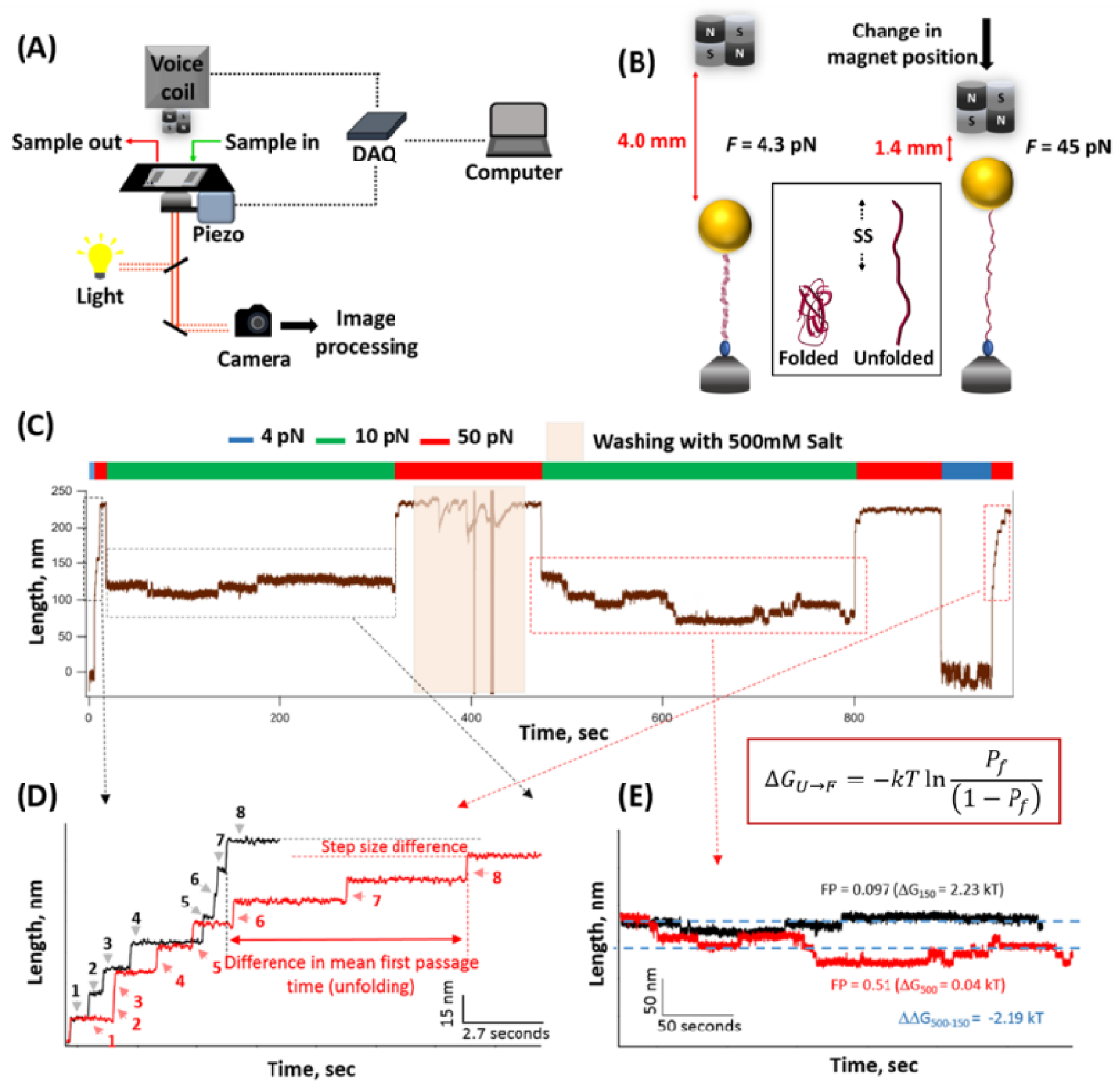
Magnetic tweezers instrumentation: **(A) Instrumentation setup:** The instrument includes a flow chamber which ensures a continue flow of the buffer containing external stimuli and a magnetic tweezers, allowing us to change the solution multiple times during experiment while tracking a single protein molecule. The applied force on protein-bound paramagnetic bead is controlled by a pair of permanent magnets placed above the flow chamber. **(B) A schematic representation of single molecule experiment:** The molecular construct comprising protein L octamer is flanked by an N-terminus HaloTag (blue sphere) for tethering to chloroalkane ligand on glass coverslip and C-terminus AviTag [HaloTag-(Protein L) _8_-AviTag] for reacting to streptavidin coated paramagnetic bead (yellow sphere). The difference between unfolded and folded length is denoted as step-size (SS) (inset in Fig. 1B). **(C) Magnetic tweezers technology to monitor multiple events of protein folding in a single molecule from a single experiment:** Firstly, the folded construct (at 4 pN) is unfolded by a constant force of 50 pN leading to eight successive unfolding steps as single molecule fingerprint. Then the force is allowed to quench at 10 pN where the fully unfolded polypeptide shows immediate elastic recoiling and folding. Then the construct is unfolded by fingerprint pulse of 50 pN and the flow cell is washed four times with 500 mM salt. As previously, the force is quenched to 10 pN for observing the equilibrium phase where any changes in folding-unfolding dynamics can be precisely traced and followed by similar high force pulse. Then the protein is completely refolded at 4 pN and unfolded again at 50 pN to check any changes in unfolding **(D) Salt increases unfolding time and promotes conformational changes:** The unfolding events at two different salt conditions are magnified and compared. At 150 mM, the octamer unfolds rapidly within 5.4 s (black trace), but at 500 mM, it takes 13.45 s (red trace). This delayed unfolding is a clear indication of higher mechanical stability of the folded protein provided by salt. Moreover, the observed difference in step size offers a hint of conformational changes. **(E) Effect of salt at low force regime:** The refolding trajectories at 150 mM (black trace) and 500 mM (red trace) are compared to investigate refolding time and folding probability. With 150 mM, the polyprotein shows folding-unfolding dynamics among 1^st^, 2^nd^ folded state whereas with 500 mM, the dynamics is markedly accelerated and attain an equilibrium phase where it hops between 4^th^, 5^th^ and 6^th^ folded states. Salt here works as a mechanical folder, allowing the protein to fold at high force

### A single real-time tracking experiment of a single protein reports salt-dependent perturbation in kinetics, thermodynamics and polymer properties of protein L

Fig. 1C demonstrates how magnetic tweezers technology measures the difference in thermodynamic and polymer properties of a single protein only from a single experiment, while introducing the salt during that experiment. In Fig. 1C, at the first phase, the folding dynamics of protein L was recorded at 150 mM salt and in the next phase, the flow chamber was washed with 500 mM salt, followed by measuring the folding dynamics of protein L in presence of 500 mM salt. In this article we used ammonium sulfate [(NH4)_2_SO_4_] as a model salt.

In Fig. 1C, first protein L is unfolded at a high fingerprint force pulse of 50 pN and subsequently quenched to equilibrium force of 10 pN to monitor folding dynamics in presence of 150 mM salt. This is followed by another high force pulse of 50 pN and washed the flow cell four times with the 500 mM (NH4)_2_SO_4_. The protein is then relaxed at 10 pN, unfolded at 50 pN, collapsed at 4 pN and again unfolded at 50 pN. Notably, the covalent bond between HaloTag enzyme and chloroalkane ligand helps the protein to remain tethered to the glass surface while washing the flow cell. Two unfolding trajectories (from 4 pN to 50 pN) at 150 mM and 500 mM salt, are magnified and compared in Fig. 1D. In 150 mM salt, the protein L construct unfolds rapidly within 5.4 s (Fig. 1D, black trace), whereas with 500 mM, the unfolding becomes retarded to 13.5 s (Fig. 1D, red trace). This result clearly demonstrates the increase in unfolding time with salt concentration due to more stable folded state. Furthermore, the same data also showed a prominent decrease in step size, which argues in favor of salt dependent conformational change in protein.

Fig. 1C are zoomed in to demonstrate the comparison between the folding probability (*FP*) and refolding kinetics, obtained from the equilibrium phase at different salt concentrations (Fig. 1E). At first, the polyprotein construct is unfolded at 45 pN to generate eight discrete unfolding steps, followed by quenching to a refolding pulse of 10 pN. During refolding, the polyprotein experiences a relaxation phase (which is non-equilibrium) and successively, an equilibrium phase where the domains exhibit folding-unfolding dynamics. The folding-unfolding transitions are observed as ascending (for unfolding) and descending (refolding) steps, suggesting a balanced condition for the substrate protein. The folding probability (FP) has been calculated by dwell-time analysis, particularly from the equilibrium phase (Supplementary table 1). Upon addition of 500 mM salt to flow cell, folding probability is altered in favor of folded state. During the first equilibrium trajectory with 150 mM salt, the polyprotein hops between fully unfolded,1^st^ and 2^nd^ folded states to give a calculated value of *FP* = 0.097 (Fig. 1E, black trace), whereas after introducing 500 mM salt, the polyprotein hops between 4^th^, 5^th^ and 6^th^ folded state by accelerating its *FP* to 0.51 (Fig. 1E, red trace). We have examined the folding probability by dwell-time analysis (Supplementary table 1). In equilibrium phase, each folded state is denoted as *I* (*I* is the number of folded states) and their population at a certain applied force is calculated as *t*=*T*_*I*_/*T*_*T*_, where *T*_*I*_ is the cumulative time spent on *I* state, and *T*_*T*_ = *∑*_*I*_ *T*_*I*_, total observation time of equilibrium phase. The FP is measured as normalized averaged state^59–61^.

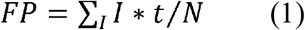

Where *N*=8 is equivalent to total numbers of domains. Detailed calculations of *FP* have been discussed in Supplementary table. 1:

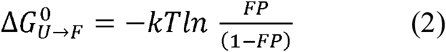

We obtained the *equilibrium* free energy difference for folding (*ΔΔG*^*0*^_*U*→*F*_) from the same experiment by analyzing the equilibrium folding under constant force of 10 pN with the help of eq. 2.^62^ We estimated the free energy in presence of 150 mM salt (*ΔG*^*0*^_*150*_) is 2.23 kT and that of 500 mM (*ΔG*^*0*^_*500*_) salt is 0.04 kT. Taken together, the *ΔΔG*^*0*^_*U*→*F*_ (*ΔG*^*0*^_*500*_ *-ΔG*^*0*^_*150*_) is around -2.2 kT which shows the folding process is more favored in presence of 500 mM salt. Moreover, from the Fig. 1E we can compare the refolding kinetics. In presence of 150 mM, two domains fold in 140 seconds but in 500 mM salt, six domains fold within the same observed time scale.

### Comparison of single experiment data with many single molecule experiments

In Fig. 2, the information obtained from the single experiment (Fig. 1C) is correlated with the results of many single molecule experiments at a constant salt concentration. In this figure many folding dynamics events of protein L were monitored at a constant salt concentration and their average results are compared with the data obtained from the single experiment (Fig. 1C).

**Figure 2:**
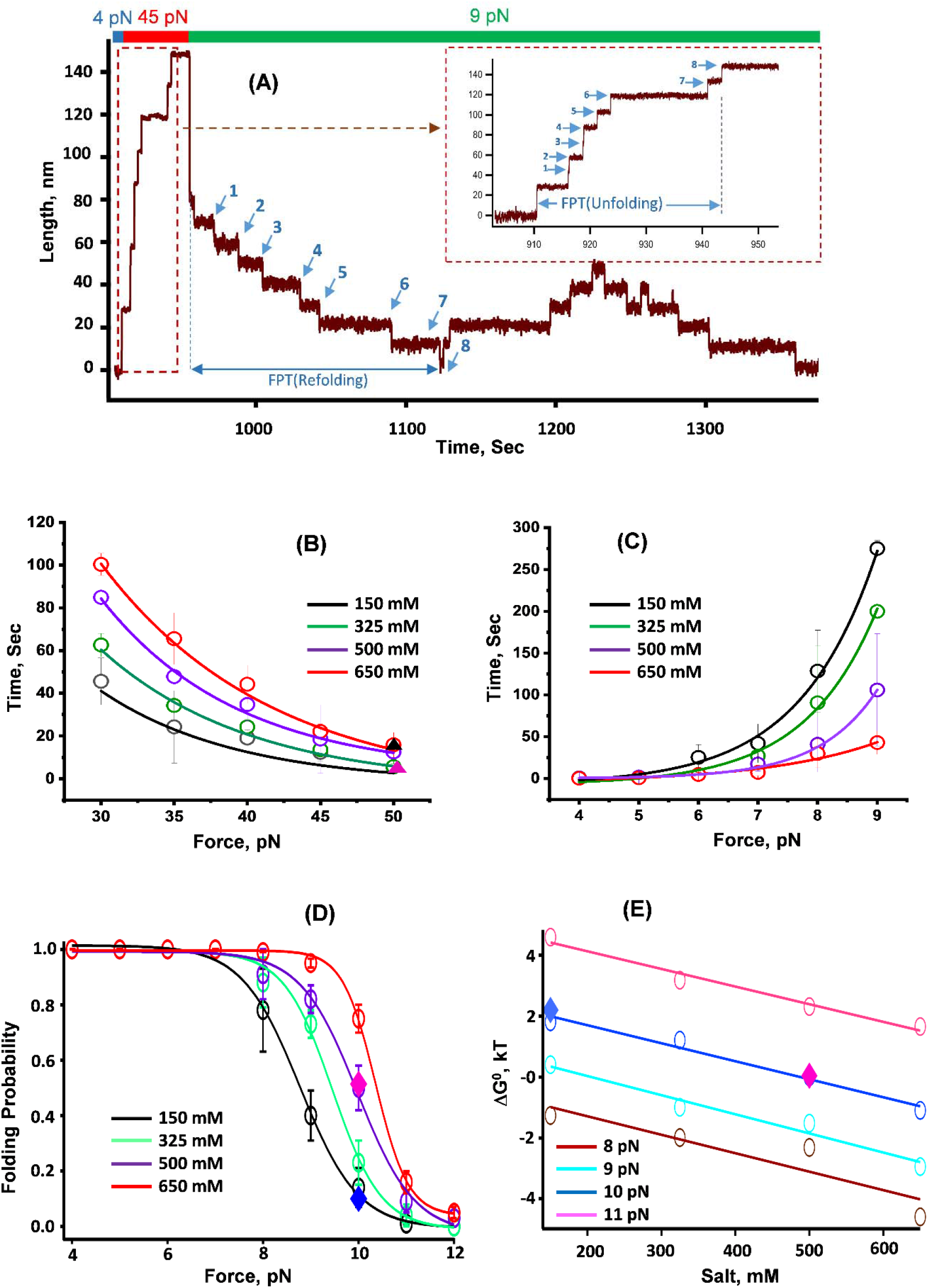
Comparison of single experiment with many single molecule experiments: **(A) Standard trajectory of protein L at 500 mM salt concentration obtained from magnetic tweezers experiment:** The protein L octamer is first unfolded at a fingerprint pulse of 45 pN leading to complete unfolding of eight domains. After unfolding, the force is quenched to 9 pN for observing the folding-unfolding dynamics at equilibrium phase. The unfolding trajectory is magnified to inset, where the first passage time of unfolding (FPT) is described as the time taken to complete unfolding from fully folded state. Similarly, FPT of refolding denotes the total time for complete refolding from fully extended state. **(B) Variation of mean-FPT for unfolding:** The MFPT of unfolding is plotted as a function of force with different salt conditions. The unfolding time decreases steadily with increasing force, however, salt, acting against to force, increases the time for complete unfolding. Notably, the data at 50 pN, acquired from single molecule analysis, shows complete unfolding with 150 mM takes 5.4 s (pink triangle), whereas it increases to 13.44 s with 500 mM (black triangle) which are consistent to the plot of unfolding-MFPT. More than eight individual trajectories are measured and averaged for each force. Error bars are standard errors of mean (s.e.m.). **(C) Variation of MFPT for refolding:** Salt reduces the total refolding time. For example, at 9 pN, in presence of 150 mM, the MFPT of refolding is 275±10 s (black trace) while with 650 mM the time decreases to 43±14 s (red trace). More than eight individual trajectories are measured and averaged for each force. Error bars are s.e.m. **(D) Folding probability as a function of salt concentration:** The folding probability as a function of force is plotted for four different salt concentrations (150, 325, 500, 650 mM). The effects of salt on folding probability increases considerably over a range 8-11 pN and attains maximal at 10 pN force. Our single experiment showed that at 10 pN, the folding probability has been increased from 0.097 (in 150 mM salt, blue diamond) to 0.51 (in 500 mM salt, pink diamond), showing a strong concentration dependent role of salt on protein folding. Data points are calculated using >2500s and over five molecules per force. Error bars represent s.e.m. **(E) Effect on** Δ**G**^**0**^_**folding**_ **with increasing salt concentration:** ΔG^0^_folding_ (kT) is plotted as a function of salt concentration at different forces ranging from 8 to 11 pN. With increasing salt concentration at a particular force, the ΔG^0^ decreases. From our single experiment data of folding probability, we calculated the ΔG^0^ at 10 pN force as 2.23 kT (150 mM, blue diamond) and 0.04 kT (500 mM, pink diamond), which are precisely coinciding with our ΔG^0^ plot.

Fig. 2A shows a standard experimental trajectory of an octamer construct of protein L at 500 mM salt concentration, where the construct is unfolded by an unfolding pulse of 45 pN force and folded by a subsequent refolding pulse of 9 pN. After applying the refolding pulse, the domains undergo an equilibrium stage, where the domains exhibit folding/unfolding dynamics as ascending/descending steps. The unfolding trajectory is magnified in the inset of Fig. 2A, where the first passage time (FPT) of unfolding is described - the minimum time taken to complete unfolding from fully folded state (time between the arrow, in Fig. 2A inset). Similarly, the FPT of refolding is calculated as time for a complete refolding from a complete unfolded state. Mean first passage time (MFPT) is determined by averaging the FPT values over several trajectories, offering a model-free metric that describes the unfolding and refolding kinetics of protein under force. Fig. 2B shows the variation of MFPT of unfolding as a function of force and salt concentration. At a particular salt concentration, the MFPT of unfolding decreases with increasing mechanical force - showing the inverse relation of protein folding with force. However, salt changes the energy landscape and unfolding time increases with salt concentration such as, at 50 pN, complete unfolding takes 5.5±2.1 s in presence of 150 mM salt whereas it increases to 12.48±1.6 with 500 mM salt. Importantly, our single molecule data is strongly correlated with the unfolding-MFPT plot such as, in presence of 150 mM salt, complete unfolding of eight domains at 50 pN takes only 5.4 s (Fig. 2B, pink triangle) but at 500 mM, the time shifts to 13.45 s, (Fig. 2B, black triangle). In contrast to MFPT of unfolding, MFPT of refolding decreases with increase in salt concentration. For example, at 9 pN force with 150 mM salt, the polyprotein completely refolds at 275±10 s (Fig. 2C, black trace) whereas in presence of 650 mM salt, complete refolding takes just 43±14 s (Fig. 2C, red trace). Thus, salt delays the unfolding kinetics and accelerates the refolding kinetics in concentration dependent manner. MFPT values has been fitted with the equation *MFPT = Ae*^*F/r*^, where *A* is a time fitting parameter and *r* is a force fitting parameter. In case of MFPT unfolding, *r* has a negative value as MFPT unfolding decreases with the applied force.

Similar to MFPT for unfolding and refolding, salt concentration greatly affects the thermodynamic energy landscape of protein folding, as observed in Fig. 1E. To thoroughly investigate that, we plot the folding probability as a function of force at different salt conditions. We observed that folding probability decreases with increasing force - showing again the inverse relation of folding and force. However, introducing salt, opposing to force, helps in protein folding and thus, escalates the folding probability. At 4-7 pN force regime, all domains fold rapidly and completely, irrespective to the salt concentration and thus, FP=1. At 12 pN, the FP decreased to zero, except 500 and 650 mM where FP are 0.03±0.02 and 0.05±0.02, respectively. However, at an intermediate force range of 8-11 pN, the folding probability is drastically raised with increasing salt (Fig. 2D). The half-point force (defined as force where FP = 0.5) shifted rightward from ∼8.8 to ∼10.4 pN at 150 mM to 650 mM range, showing a higher intrinsic ability of protein to remain folded against the force in presence of salt (Supplementary Fig. 1 to Supplementary Fig. 4). Fig. 2E shows a comparative data of free energy difference (*ΔG*^*0*^) within a span of 8-11 pN force, where both folding-unfolding transitions occur within experimental time-scale and *ΔG*^*0*^_*U*→*F*_ was calculated from eq. 2. Our result shows salt favors folding and thus, *ΔΔG*^*0*^_*U*→*F*_ decreases with gradual increment in salt concentration and the result is consistent for all the force range from 8-11 pN. Indeed, these results are in agreement with our single experiment data shown in blue (150 mM) and pink (500 mM) diamond in Fig 2D and 2E.

### Effect of salt on the free energy landscape of protein folding

To determine how salt reshapes the free energy landscape of protein folding, we investigated the unfolding rates of protein L at different salt conditions under 30-45 pN force range. By fitting single exponential to the traces plotted, we defined the observed unfolding rate as *k*_*u*_*=1/*τ at different salt conditions, where τ is the time constant measured from the exponential fits. In Fig. 3A, we have plotted representative unfolding traces in the presence of 650 mM salt concentration at 40 and 45 pN force, showing the effect of force on proteinL unfolding. For the measured values of *k*_*u*_ within 30 to 45 pN force regime, a linear relationship between *lnk*_*u*_ and force has been observed for different salt concentrations (Fig. 3B) and expressed as a Bell-like (Bell et al., Science, 1978) relationship^63^ (eq. 3).

**Figure 3:**
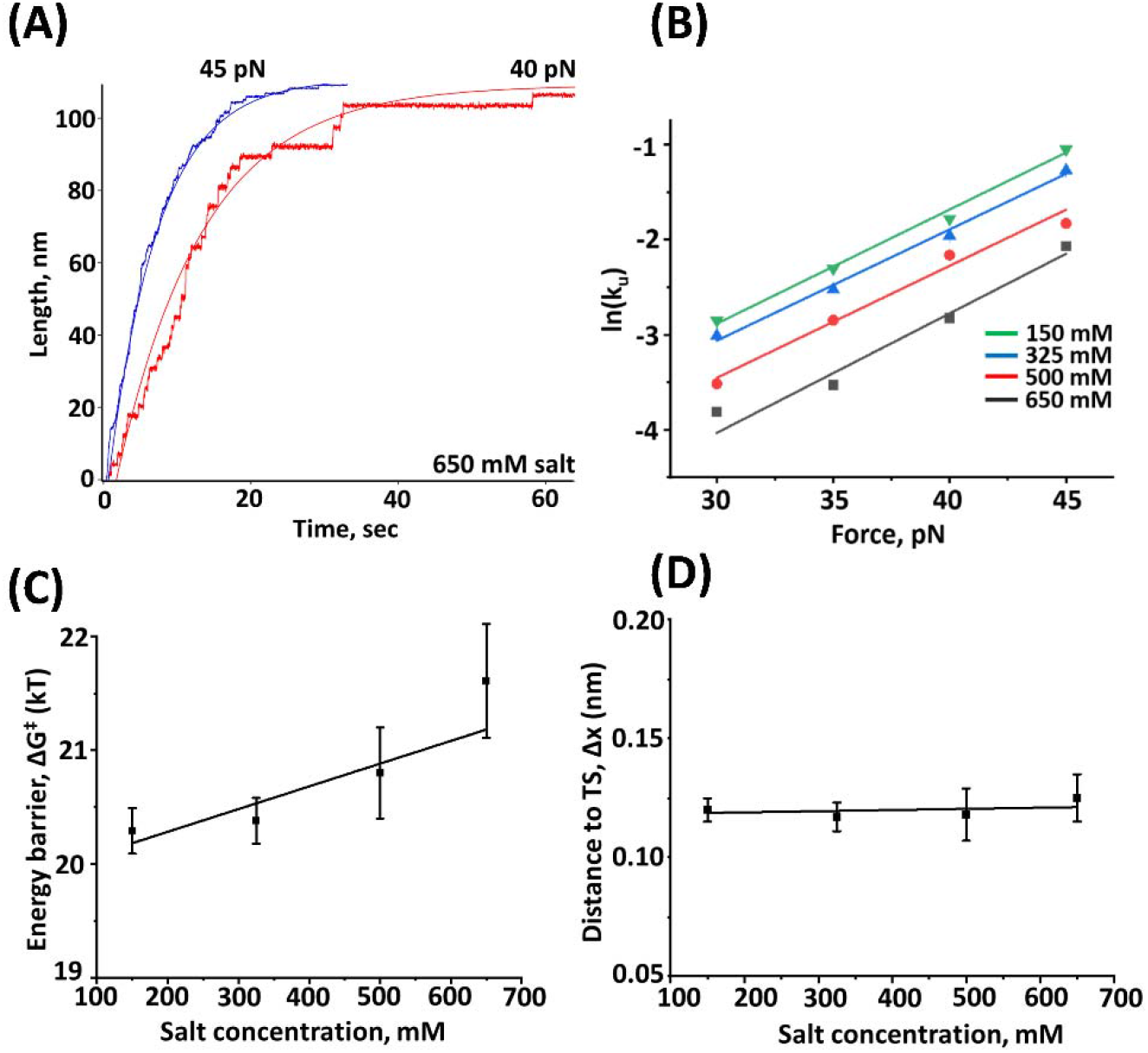
Variation in the properties of free energy barrier under different salt concentrations: (A) Sets of unfolding trace averages are plotted at 650 mM salt concentration with 40 pN (red trace) and 45 pN (blue trace) force. Single-exponential fits (continuous lines with different traces) measure the time constant, τ of the protein L unfolding. (B) The unfolding kinetics k_u_=1/_τ_, as a function of force has been plotted at different salt concentrations (150 mM, green; 325 mM, blue; 500 mM, red; 650 mM, black). A linear relationship of ln k_u_ with applied force has been observed for various salt concentration, following a bell-like equation. Each data points of unfolding rates are calculated from more than 20 individual unfolding traces. Error bars are very small that lies within the data points. (C) Salt increases the height of the free energy barrier (ΔG^†^) while increasing the concentration from 150 mM to 650 mM. Error bars are standard errors. (D) Salt does not change the distance to the transition state (TS). Error bars are standard errors.

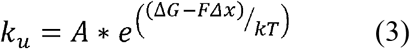

In this equation, *A* is the pre-exponential factor with the observed value of 10^6^ s^-1^, *ΔG*^*†*^ is free energy barrier in absence of force; *F* is applied force that reduces the free energy barrier by *FΔx* amount where *Δx* is distance to the transition state (nm), along the reaction coordinate^64,65^. We have observed that energy barrier height increases from 20.3±0.2 to 21.6±0.5 kT (Fig. 3C) with the increasing salt concentration while not affecting the distance to the transition state (Fig. 3D). Notably, similar calculations for the refolding rates could not be described by the Bell-like equation as their folding rate declines sharply under the low force regime (4-10 pN). This nature of dynamic folding transition is strongly attributed to the varying entropic elasticity of the extended polyprotein under different forces^66–69^. Thus, the refolding transition state distance of protein L under force, is uneven due to substantial shift of the unfolded state positions which become further complicated at varying ionic concentrations.

### Conformational compaction of protein described by ‘electrolyte FJC model’

Although salt effects on protein folding is thoroughly characterized, how electrolytes influence conformational flexibility under force is not known. To systematically investigate that, we monitored the salt dependent conformational changes of a single protein under a wide force range. With increasing salt concentration at a particular force, we observed a gradual decrease in step sizes (Fig. 5). We determined the Kuhn length (Supplementary Table. 2) and Kuhn segment (Table 1) for protein L at different salt concentration by plotting the step size data to freely jointed chain (FJC) model (eq. 4) against different forces at constant salt concentration (Fig. 4A-4D)^66,70^. We selected FJC model over the WLC model in our magnetic tweezers, since WLC model calculates the force from the given extension, while FJC model allows us to determine the extension from the applied force.^71^ Recently, it has been reported that despite of similar derivation of force-extension relationship by WLC model, FJC model is analytically more convenient due to an explicit force dependency of step size.^72^ The protein is considered as linear single stranded polymer having maximum extended length (*L*_*c*_, contour length) with *N* numbers of uncorrelated Kuhn segments of length *L*_*k*_ (Kuhn length) where, *L*_*c*_*= N * L*_*k*_ and each of those segments can take any 3D orientation. Contour length of protein L is independent to salt concentration (Fig. 4E) and is similar to the value that we observed in previous force spectroscopy studies^60^, while the Kuhn length decreases with salt concentrations (Fig. 4F; Supplementary Table 2). *N* is also calculated and shown in Table 1.

**Table 1:** Variation in Kuhn segments with increasing salt concentration

| Salt Concentration (mM) | Kuhn segments (N) |
| --- | --- |
| 150 | 14.1±0.2 |
| 325 | 15.8±0.1 |
| 500 | 20.2±0.1 |
| 650 | 28.6±0.1 |

**Figure 4:**
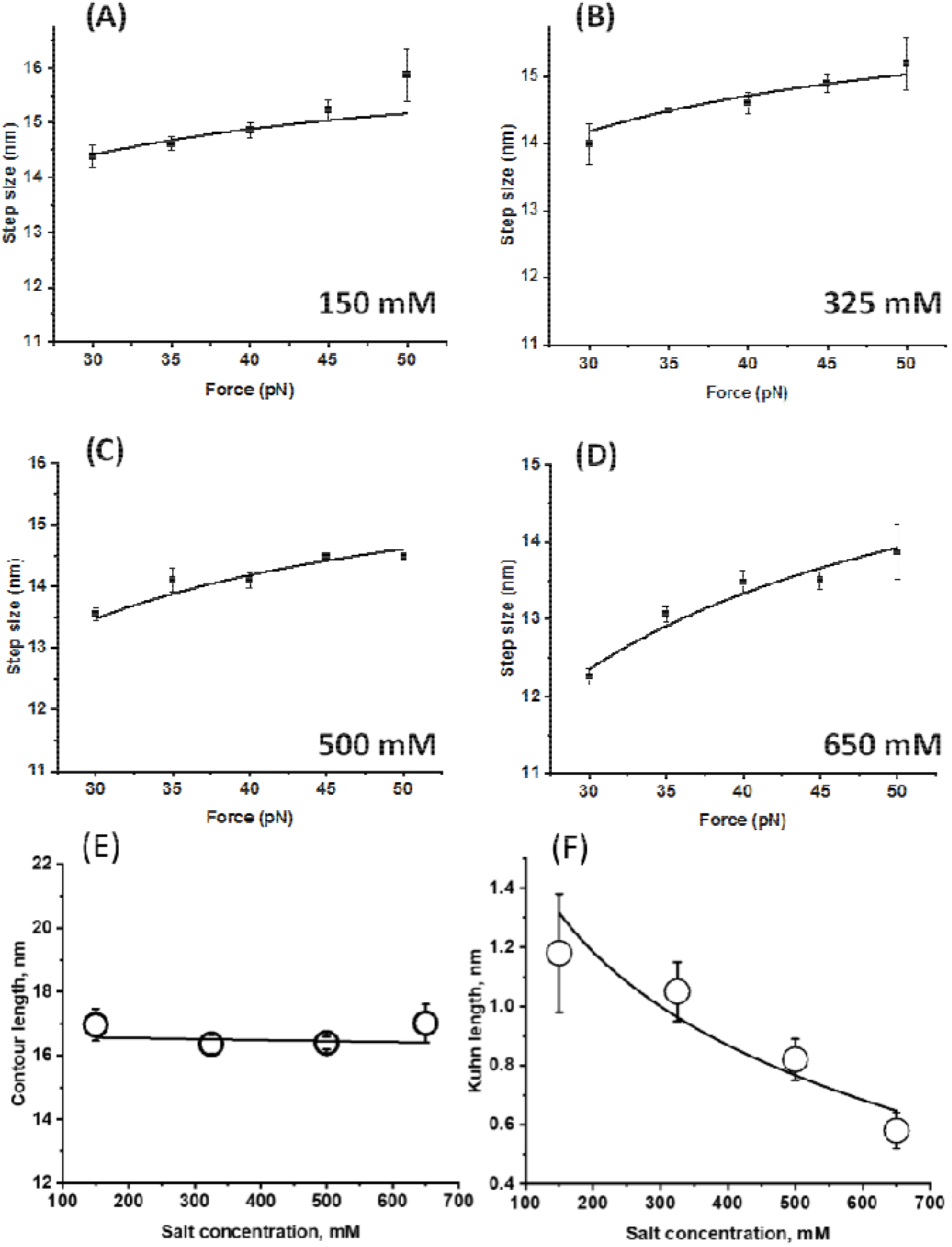
Variation in Contour length and Kuhn length of protein L at varying salt concentrations: **(A-D) Variation in the step size of protein L at varying forces with different salt concentration:** Step sizes are plotted as a function of force at different salt concentration and fitted to the simple FJC model. We observed that the step size decreases with the salt concentration and the associated Kuhn length has also been observed to decrease from 1.18±0.2 to 0.58±0.1 nm (Supplementary Table 2). More than 50 individual molecules were tested and error bars are s.e.m. **(E) Contour length:** Salt does not change the contour length of the protein L. More than 5 individual molecules were tested and error bars are standard errors. **(F) Kuhn length:** Salt decreases the Kuhn length while increasing the concentration from 150 mM to 650 mM. More than 5 individual molecules were tested and error bars are standard errors.

**Figure 5:**
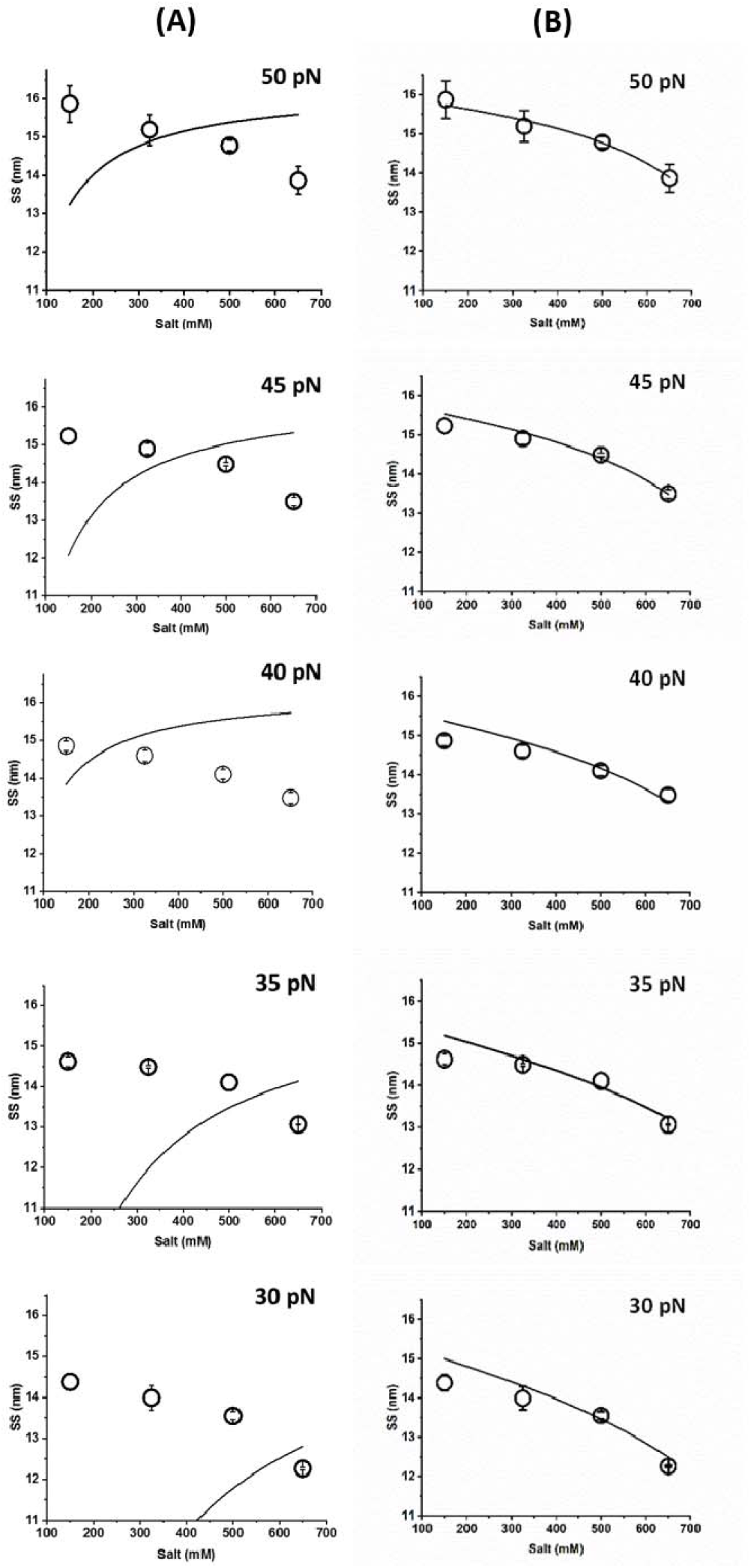
Comparison of FJC and Ec-FJC model for fitting the step size data against varying salt concentration: **(A) Simple FJC model:** The step size data against the varying salt concentration are not fitted to the model due to the absence of a salt concentration term and only explicit force dependency of step size data. However, while fitting to (B) **Ec-FJC model:** The step-size are plotted as a function of varying salt concentration at different constant forces and fitted to the electrolyte-FJC (Ec-FJC) model. We observed that these data are fitted well to the Ec-FJC model due to the incorporation of salt concentration term in the model. More than five individual trajectories are measured and averaged for step size at different salt concentration. Error bars are s.e.m.

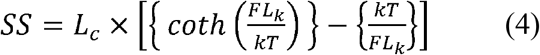

Where, *SS* is step size or extension; *F* is force (pN); *kT* is thermal energy and has a value of 4.11 pN.nm and *N* is the number of uncorrelated Kuhn segments of length *L*_*k*_ where, *L*_*c*_*= N * L*_*k*_ and each of those segments can take any 3D orientation.

We plotted protein L step sizes with different salt concentration (150-650 mM) at a particular force and fitted to simple FJC model (eq. 4). Since the simple FJC model does not include the salt concentration term and only describes the force-dependent nature of step size, it could not fit to the step size data against the varying salt concentration (Fig. 5, A panel). To investigate the effect of salt concentration on protein extension or step size under a particular force, we observed a log-dependent Kuhn length change with salt concentration ^27,73^ and explained as: 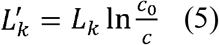, where 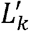 is modified Kuhn length due to electrolyte concentration change and *L*_*k*_ is Kuhn length of the polypeptide chain.

This modifies the FJC model to electrolyte FJC (Ec-FJC) to determine the step size value as a function of salt concentration, which fits the step size data at different salt concentration (Fig. 5, B panel). C is the concentration of the salt and *c*_*0*_ is a salt concentration fitting parameter, used to make the term 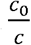 unitless.

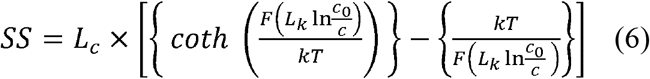

Here *L*_*k*_ value and its relation work in the range of 150-650 mM ammonium sulfate salt concentration. The relation between step size and salt concentration could be more complex and we found the equation works better in the range of 325-650 mM salt concentration. Therefore, the salt concentration term of Ec-FJC model allows it to signify the phenomenological expression of electrolyte dependent behaviour of protein step size at a particular force with better fitting.

### Salt irrespective nature of conformational changes in proteins

To cross-check whether the conformational changes are specific to ammonium sulfate salt, we repeated all the experiments of protein L with physiological sodium chloride (NaCl) salt and observed similar result to that of ammonium sulfate. NaCl increases folding probability from 0.14±0.06 to 0.28±0.03 at 10 pN (Supplementary Fig. 6), by compacting the step size of protein L in a concentration dependent manner (Supplementary Fig. 7).

To reconcile our observation, we further extended our study to another well-characterized force dependent protein talin R3-IVVI, which has been reported to have higher mechanical stability than the wild type R3 domain^55–58^. We have investigated the force-dependent folding dynamics of R3-IVVI domain in both 140 and 1000 mM NaCl. At 140 mM NaCl, the domain stays mostly in the folded state at 8.5 pN, unfolded state at 11.5 pN and occupies both 50% folded and unfolded state at 10 pN (Fig. 6A) while the folding dynamics shifts towards higher force regime in presence of higher NaCl concentration. For example, with 1000 mM NaCl, the protein stays mostly in the unfolded state at 14.5 pN, folded state at 11.5 pN and at 13 pN, it is populated in almost 50% folded and 50% unfolded state (Fig. 6B). Interestingly, we found a compaction in step size, similar to protein L, at higher concentration of NaCl (histograms of Fig. 6A and 6B). We found a compaction from 19 nm to 16 nm in presence of 140 mM and 1000 mM NaCl, respectively (Fig. 6D). Similar to step size, the half-point force has also been observed to shift to high force while increasing the salt concentration (Fig. 6C). These results generalize our hypothesis of salt-dependent compaction, in correlation with folding.

**Figure 6:**
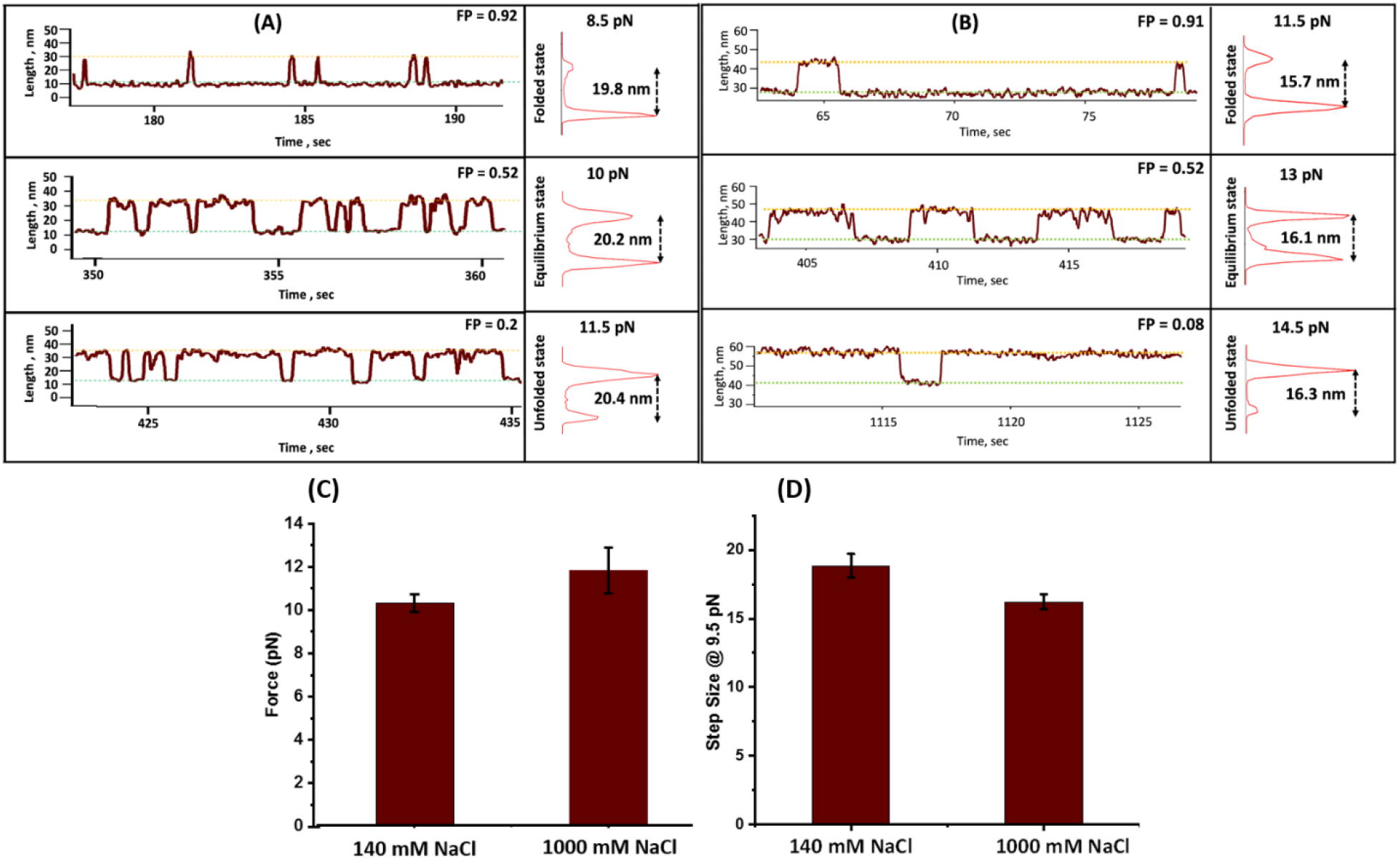
Effect of NaCl on folding dynamics of talin: **(A) Representative traces of talin R3-IVVI at different forces with 140 mM salt concentration:** folding dynamics of the R3-IVVI domain is strongly force-dependent. At 8.5 pN force, it is mostly in folded state with the folding probability of 0.92 (top trace), whereas at 11.5 pN, it is mostly unfolded with the folding probability 0.2 (bottom trace). At 10 pN, folding probability is 0.52, where both the states present in almost equal population. **(B) Representative traces with 1000 mM salt concentration:** The folding dynamics of the domain has been observed to shift towards higher force regime in presence of 1000 mM NaCl. At 11.5 pN, the domain mostly stays in folded state with the folding probability of 0.91 (top trace), whereas at 14.5 pN, it is mostly unfolded with a folding probability of 0.08 (bottom trace). At 13 pN, the folding probability is 0.52, where both the states present in almost in equal population. **(C) Effect of NaCl on half-point force of talin:** At 140 mM NaCl, the half-point force is 10.03±0.45 pN while in the presence of 1000 mM NaCl, it increases to 11.5±1.03 pN. Each data bars are calculated from the averaged half-point force of six individual molecules. Error bars represent s.e.m. **(D) Effect on Step size:** At a constant force of 9.5 pN, the step size is 18.84±0.86 nm with 140 mM NaCl, which decreases to 16.22±0.53 nm in the presence of 1000 mM NaCl. Each data bars are calculated from the averaged step size of six individual molecules. Error bars represent s.e.m.

## Discussion

Force spectroscopy studies revealed that salt ions of Hofmeister series influence the stability of substrate proteins under force, by modulating their thermodynamic parameters and screening electrostatic interactions. For example, sulfate ions have been observed to decrease the mechanical stability of titin domain by ionic screening of electrostatic interactions and similarly, borate and perchlorate reshape the free energy landscape by shifting the transition state towards the denatured state.^33^ However, how varying ionic strength influence the multiple physical properties such as, *ΔG*^*0*^, kinetics and polymer elasticity of a single protein molecule under mechanical constrains is not studied. Notably, correlation among these physical properties was also missing in the literature which can illustrate a comprehensive picture of protein chain physics under varying ionic strength. The single molecule analysis, performed by magnetic tweezers technology, reveals an integrated approach of studying protein folding by combining kinetics and thermodynamics to polymer physics from a single experiment (Fig. 7). Here, we monitored both mechanical unfolding and refolding at varying salt concentrations in a wide range of mechanical force (Fig. 7A). Our result shows, salt escalates the conformational flexibility by increasing the number of Kuhn segments, promoting the higher degrees of native contacts to favor the folding (Fig. 7B and 7C), which also reflects in the accelerated refolding kinetics and delayed unfolding kinetics. Interestingly, from the kinetics data, we observed how salt changes the energy landscape by increasing the height of the free energy barrier with respect to the folded state, keeping the distance to the transition state constant. These findings are in agreement with a study in which sulfate ions stabilizes the folded conformation of protein L domain by increasing the surface tension of the solution.^32^ Here we used ammonium sulfate solution which as a poor solvent, collapses the polypeptide in a concentration dependent manner due to an enhanced monomer-monomer interaction and a diminished monomer-solvent interaction. This collapse the polypeptide chain, forming a more compact globule state of the protein substrate. This salt-induced collapse transition could occur simultaneously with the folding transition; however, the pathway could be distinct from that of folding. From a folding landscape perspective, the conformational entropy could be reduced in the collapsed state and thus, their stability is lowered than the folded state. This entropic loss may enhance the protein folding^74^. Due to the globular folded conformation, ammonium sulfate may collapse the protein L polypeptide by modulating their hydrophobic interactions which eventually results in the clustering of non-polar residues. This collapse transition is mainly driven by the formation of van der Waal’s interaction which are non-specific and presumed to be very rapid^75,76^. However, it is suggested that collapse formation is not only attributed to the hydrophobic interactions but also to hydrogen bonding in the polypeptide backbone which is either non-specific (long range) or specific (short range)^77–80^. Our work is coinciding with simulation study where it has been shown that poor solvents redesign the Ramachandran plot by redistributing polypeptide dihedral angles φ and Ψ to largely attribute in the fold topology during the early collapse in the folding pathway. Indeed, the ‘bridge region’ between the β-basin and α region in Ramachandran plot is not allowed in good solvents while in poor solvent the bridge region is favored and thus, induces the folding during molecular compaction of denatured state ensemble^81–83^. The resulting molecular compaction facilitates the protein folding by decreasing both their conformational entropy and conformational spaces, screened during folding.

**Figure 7:**
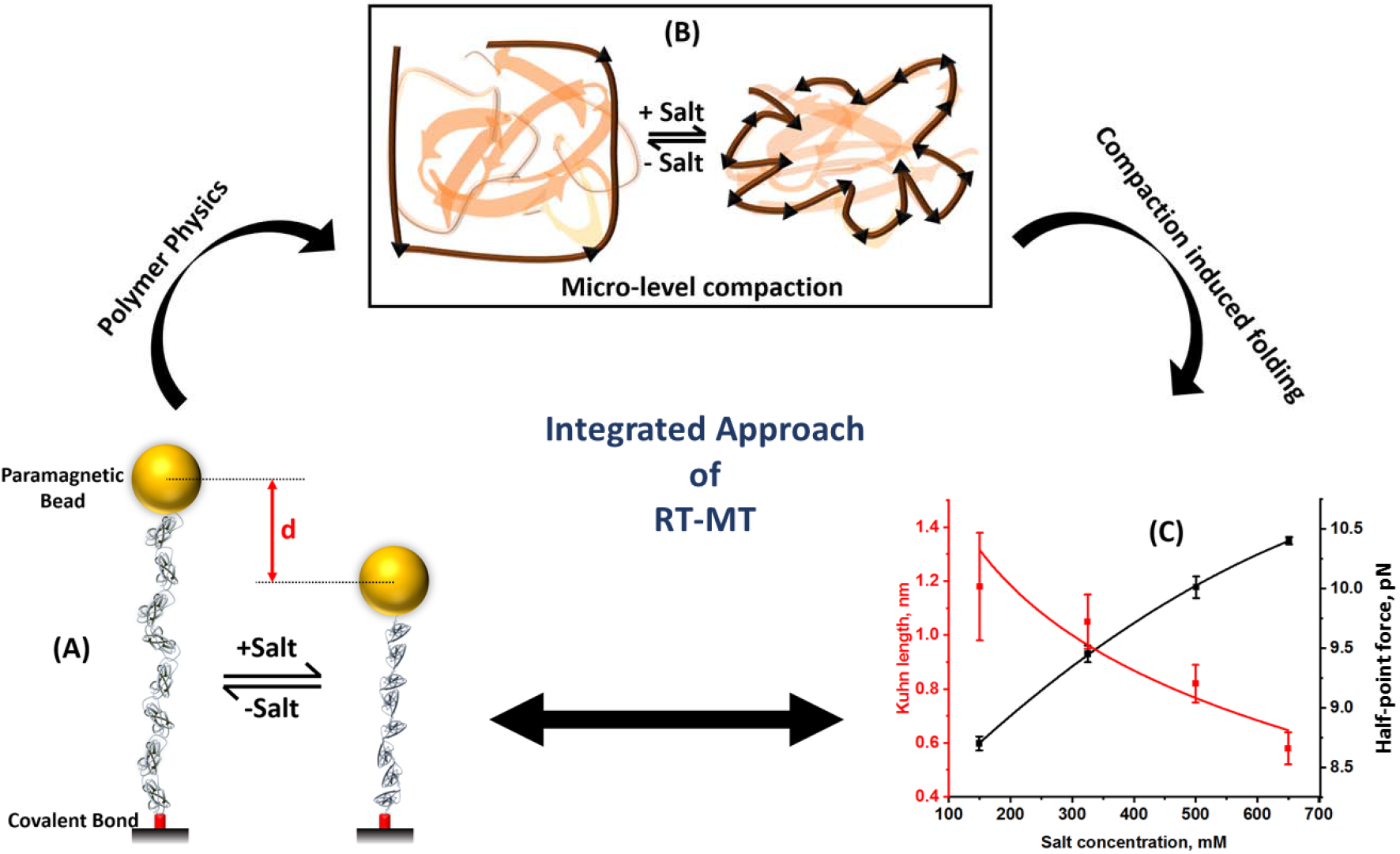
Magnetic tweezers technology has a multifaceted application to integrate polymer physics, kinetics and thermodynamics from a single experiment: **(A)** Salt induces the conformational changes in protein domains that observed from the decrease in step-size, ‘d’. **(B) Salt induced change in polymer property:** Addition of salt causes compaction of polypeptide chain. This compaction can be analyzed by illustrating the micro-level compaction, where salt increases the number of Kuhn segment in protein polymer to promote flexibility. **(C) Molecular compaction facilitates protein folding:** Kuhn length (Supplementary Table 2) and half-point of folding probability (black square, Figure. 2D) are plotted as functions of salt concentration. Salt decreases the Kuhn length of the protein to induce the mechanical flexibility while simultaneously increases the half-point of folding probability promoting a higher intrinsic ability to remain in the folded state. Thus, higher order of native contacts and successive molecular compaction, due to increased mechanical flexibility, facilitates the stability of the folded state.

Different models of polymer compaction including, mean-field and scaling theories have been proposed to understand the microscopic chain physics of proteins. All of these models correlate the dimension and length of the chain with their free energy landscape. The dimensions of both intrinsically disordered proteins (IDPs) such as, prothymosin α, HIV integrase and foldable proteins (such as, cold shock protein, cyclophilin A, spectrin domain R15 etc.) are strongly dependent on solvent quality and indeed, these have been extensively studied by single molecule technologies^84,85^. The molecular compaction of polypeptides, at both their unfolded and folded state, are defined as radius of gyration and critically determined by the solvent quality which can be defined by the empirical relationship^86^

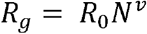

where *R*_*g*_ is radius of gyration, *R*_*0*_ is a prefactor, *N* is no of Kuhn segments and *v* is defined as scaling exponent-a measure of the conformational fluctuations of polypeptide under a solvent condition. Scaling exponent of polypeptide chains largely depends on the solvent quality and has a value of 0.6≥*v*≥0.33. In good solvent, polypeptide chain forms extended state by enhancing the residue-solvent interactions than intra-residue interactions, giving a value of *v*∼0.6, while it reduces to only 0.33 in the presence of poor solvent such as ammonium sulfate, by maximizing the intra-residue contacts^87^. Additionally, prefactor *R*_*0*_ is a multifactorial parameter which is a function of excluded volume, chain thickness, and Kuhn length^88^. Interestingly, from our observation, we can predict that ammonium sulfate facilitates the collapse transition, possibly by affecting the Kuhn length and therefore, the *R*_*0*_. Interestingly, water acting as poor solvent for the polypeptide, decreases the *v* = 0.46 for the unfolded state of proteins which is less than 0.58 but higher than 0.33, suggesting the compaction of the unfolded states. At molecular scale, different mechanisms have been suggested for the salt-protein interactions due to dispersed charge seperation and distribution of polar and non-polar residues which further complicates the effect of different salts on protein stability under force. Kosmotropic ions such as, sulfate, phosphate and fluorides stabilize the folded state of protein which are evident from the positive slope in melting temperature of protein with the salt concentrations. By contrast, denaturant activity of different salts originates due to weak nonspecific interaction between salt ion and polar region of polypeptide, especially in the polypeptide backbone.^32^ In heteropolymer theory, the resultant intra-chain interaction energy can be determined as the sum of two mean-field terms: backbone interactions and side-chain interactions. This intriguing ability of poor solvents originates from the attractive backbone interactions in the polypeptide. Similarly, ammonium sulphate salt as poor solvent thought to reduce the scaling exponent of protein L in the intermediate range (0.58 > *v* > 0.33) that further reduces the radius of gyration of the unfolded states. Furthermore, molecular simulation of protein L in water medium suggested the compaction of unfolded state^89,90^. The unfolded state of protein L molecule undergoes rapid compaction in native condition with ∼10% larger than its folded state which apparently suggests the compaction of unfolded state^91^. Although the salt mediated compaction is still highly debated with possible attribution from either unfolded state compaction or that of folded state or even from both the states, chain compaction into a very dense state diminishes the diffusive chain fluctuations by inducing internal friction that increases with elevated compaction of unfolded proteins^92,84,85,93–95^. Additionally, in poor solvent, the protein behaves like heteropolymer, resulting in an initial collapse followed by secondary structure formation mainly arisen due to specific contacts which in turn greatly speed up the folding. Similarly, we observed ionic strength favors protein folding phenomena by concomitantly increasing the native contacts during structure formation and facilitating the thermodynamics of folding (Fig. 7C) which strongly reconciles the compaction driven protein folding.

In cells, ion homeostasis is pivotal to maintain cellular functions by constantly changing the ionic environment that affect the elasticity of different biological polymers such as, DNA, RNA or proteins. Although polymer properties of nucleic acids and polysaccharide, being intricate polymer with large persistence length and charged backbone, have been well studied^12,27,28,96–103^, such behavior of proteins has not been reported in detail, especially how the ion driven conformational flexibility of proteins changes under force. Previous measurement of titin PEVK domain using AFM have not provided a consensus picture of ion-dependent molecular elasticity as the study used a random coil domain which has been reported to exhibit distinct structural and mechanical properties than globular Ig and Fn domain of titin^17,104^. Additionally, the measurement of polymer scaling behaviour in both intrinsically disordered and random coil proteins are not in agreement with that of structured protein and thus, a comprehensive picture of molecular chain elasticity in structured protein is required. To obtain deeper insights into this, we characterize the protein L as a polymer where their step-sizes are described by the simple FJC model. Since, the *L*_*c*_ and *L*_*k*_ are known to be fixed for a protein polymer in a given ionic solution; the existing FJC model only describes the step-sizes as a function of force^66,70^. We have modified the FJC model to analyze the conformational compaction of protein polymer in varying ionic environment. We observed that *L*_*c*_ of protein L is independent to salt concentration, while the *L*_*k*_ decreases with the salt concentration. This is also relevant to the nucleic acid, which exhibits very weak dependence to the ionic strength^20^. Indeed, the mechanosensing polyprotein like microtubules and actin have relatively higher persistence length (*L*_*p*_) and thus show higher bending rigidity that allows them to form crosslinking structures^105,106^ in cell cortex and protrusion. Moreover, due to extended topology they are constantly exposed to the varying ionic environments that change unfolding pattern of their domains during interaction with other proteins that further can decode unprecedented physical viewpoints of cell division and structure maintenance. Interestingly, post-translational modifications in polypeptides change their resultant charge distribution that alters their elasticity and end-to-end distance.^107^ In physical perspective, the electrostatic interaction among charges in a polypeptide is regulated by ionic strength which effectively alters their electrostatic contribution to *L*_*k*_ or *L*_*p*_ by changing Debye screening length (λ_*D*_). Interestingly, the cellular compartments involved in secretory pathway have a well-defined pH continuum from the neutral pH to acidic pH that could affects the folding and stability of secreting protein as well as resident proteins. Stability of bi-histidine mutant of secretory protein GB1 decreases under force, upon reducing the pH from 8.5 to 3 due to electrostatic repulsion at acidic pH which in turn also affect their elastic properties.^29^ Illustrating their altered polymer properties in dynamic ionic environment could reveal new horizon for studying protein functions.

### Extraction of multiple physical properties from a single experiment and comparison with many single molecule experiments

Comparative analysis of molecular properties, before and after interaction, is appropriate if they could be performed by a single technique - as every methodology requires different chemical or physical manipulations, which introduce associated artifacts. The comparison could be even more precise if carried out on the same molecule. From a single experiment on the exact same protein individual, we are not only able to measure different molecular properties but also quantify their ionic strength dependent changes by re-measuring the exact same protein specimen under varying salt condition. We observed that protein L unfolds fast in the presence of low salt condition, while that very same molecule exhibits slower unfolding kinetics upon the addition of higher salt concentration (Fig. 1D). By contrast, the refolding kinetics has been observed to increase in the exact same protein individual in the presence of higher salt concentration. At 150 mM salt concentration, protein L construct is only able to refold its two domains within 140 s; while in the presence of higher salt concentration, it can refold upto six domains, increasing its refolding efficiency under equilibrium force condition (Fig. 1E). Notably, we compared the kinetics data only from a single experiment to many single-molecule experiments (Fig. 2), performed on different protein individuals at varying salt concentration under force; and observed that both the data are in close agreement (Fig. 2B and 2C). Additionally, we monitored how salt modulate folding dynamics of substrate protein and presented the data as FP under two different salt conditions (Fig. 1E) at equilibrium force of 10 pN. We found that protein L construct exhibit hopping between fully unfolded, 1^st^ and 2^nd^ folded states at 150 mM salt concentration, giving a lower FP; while the very same molecule upshifts its dynamics between 4^th^, 5^th^ and 6^th^ states after introducing 500 mM salt, escalating the FP to higher value (Fig. 1E). Similarly, while comparing the data extracted from this single experiment to many experiments, FP data has been found to be strongly correlated with each other (Fig. 2D). From the electrolyte-modulated folding dynamics in single protein individual, we measured the free energy difference of protein folding from the FP values by eq. 2 (Supplementary Figure 5). We observed that free energy difference of a decreases from 2.23 to 0.04 kT while increasing the electrolyte concentration from 150 to 500 mM (Fig. 1E).

The conformational change of a protein polymer under varying ionic condition is difficult to measure on the very same protein individual. Here, we observed a significant decrease in the step size of protein L domains from 15.4 nm to 15.1 nm upon increasing the salt concentration from 150 to 500 mM (Fig. 1D). Fitting these data to canonical FJC model results in the decrease of Kuhn length from ∼1.3 to ∼1.02 nm, which is also within the range of Kuhn length, described in the Fig. 4F. This certainly increases the Kuhn segments in protein L polypeptide, signifying the generation of more collapsed state of protein L due to the salt effect which facilitates the native contacts of the polypeptide to accelerate the protein folding under force. Therefore, only a single experiment allows us to extract multiple physical properties of a protein molecule, as well as their electrolyte-dependent change under varying condition. This approach quantifies both the elastic properties and unfolding-refolding dynamics, simultaneously from a single experiment. By correlating both these molecular properties from the one single experiment, we directly observe that conformational compactness of a protein modifies its energy landscape.

## Acknowledgement

We thank Ashoka University for support and funding. S.H. thanks DBT Ramalingaswami Fellowship and DST SERB Core Research Grant for funding. We sincerely appreciate Prof. Julio Fernandez (Columbia University) for helping us with the magnetic tweezers set up. We thank Dr. Rafael Tapia-Rojo (Columbia University), Dr. Edward C Eckels (Columbia University), Prof. Somendra M. Bhattacharjee (Ashoka University) and Prof. Gautam Menon (Ashoka University) for the critical analysis of the manuscript.

## Supplementary information

### Material and methods

#### Plasmid construct and protein Expression

Polyprotein constructs of protein L and talin were engineered using *Bam*HI, *Bgl*II and *Kpn*I restriction sites in pFN18a expression vector^1,2^. The protein L construct contains eight Protein L B1 domains and talin construct is comprised of a single R3-IVVI domain, followed by eight tandem I91 repeats. Both the constructs are flanked by an N-terminal Halo-Tag enzyme and a C-terminal AviTag for biotinylation. For the purification, a His_6_-tag is also present immediate before the AviTag sequence^3,4^. For protein expression, *Escherichia coli* BL21 (DE3) competent cell were grown in LB media with 50 μg/ml carbenicillin antibiotics at 37□ C until optical density (OD) at 600 nm reached 0.6-0.8. Further, the protein overexpression was carried out by inducing with 1 mM Isopropyl β-D-1-thiogalactopyranoside and, followed by overnight incubation at 25□ C.

Detailed sequence of the construct is listed below with colors labelled for corresponding component in the construct. There is a short dipeptide linker between two adjacent components of the construct are underlined.

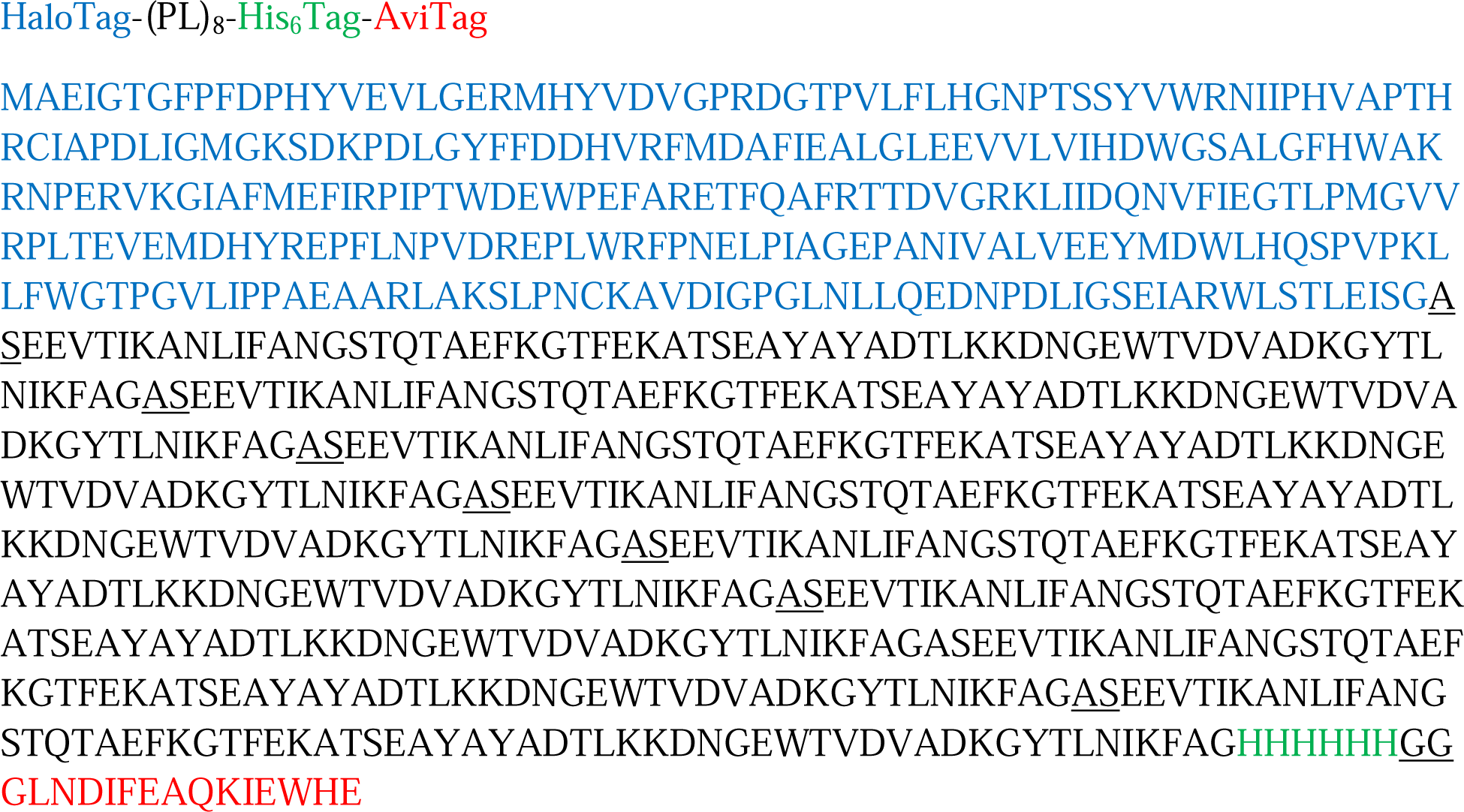

#### Protein Purification

After inducing the protein overexpression for overnight, the cells were centrifuged at 8000 rpm for 10 minutes and subsequently harvested in 50 mM sodium phosphate buffer pH 7.4 with 300 mM NaCl and 10% Glycerol. Lysozyme and 10μl Phenylmethylsulfonyl fluoride were used as protease inhibitors. The dissolved pellet was incubated in ice for 20 minutes and shaken continuously on rocking platform at 4□C for 10 minutes. Then Triton-X, DNase, RNase and MgCl_2_ were added subsequently and again put in rocking platform for 10 minutes at 4□ C and the cell were homogenized using French press at 19 psi. The lysate was centrifuged at 4□ C and supernatant is collected. Proteins were purified from lysate using Ni^2+^-NTA affinity chromatography column of ÄKTA™ Pure (GE healthcare). The sodium-phosphate buffer with 20 mM imidazole is used as binding buffer to bind the protein in column which in turn was eluted by using the same buffer with 250 mM imidazole. After purification, the polyprotein was biotinylated by the biotinylation kit (GeneCopoeia). The reaction mixture containing Protein L was kept overnight at 4□C followed by further purification by size exclusion chromatography with the Superdex-200 HR column of GE healthcare.

#### Real time magnetic tweezers methodology

Real time magnetic tweezers (RT-MT) was built on an inverted microscope using an oil-immersion objective, attached with a nanofocusing piezo. The chamber was flashed using a collimated white LED light and the diffraction pattern images were collected with ximea camera (MQ013MG-ON). Force is applied using a pair of neodymium magnets to paramagnetic beads (Dynabeads M-270) attached with the protein. A linear voice coil, placed just above the sample chamber, was used to control the position of the magnets. We use multifunctional DAQ card for controlling the voice coil and piezo actuator and for getting accurate data.

The experiments are performed with glass chamber made by two sandwiched glass slides with the advantage of inlets and outlets. At first, glass surfaces were cleaned by Hellmanex and ethanol. Then they are silanized by incubating for 20 minutes in 1% solution of (3-Aminopropyl) trimethoxysilane (sigma Aldrich, 281778) in ethanol. The fluid chambers were then functionalized with glutaraldehyde (Sigma Aldrich, G7776) and HaloTag amine (O_4_) ligand (Promega, P6741). The non-magnetic beads (2.5-2.9 μm, Spherotech, AP-25-10) were added as a reference bead. Finally, to block nonspecific interactions, the chambers were washed with freshly prepared blocking buffer (20 mM Tris-HCl pH 7.4, 150 mM NaCl, 2 mM MgCl_2_, 1% BSA, 0.03 % NaN_3_) and left them to passivate at least for 5 hours at room temperature. Single molecule experiments were carried out with 1-10 nM protein L solution in 20 mM Tris buffer with varying ammonium sulfate concentration^5^. Briefly, we separately prepared two buffers: 1) 20 mM Tris buffer, containing only 10 mM ascorbic acid of pH 7.2 and 2) 20 mM Tris buffer containing 10 mM ascorbic acid, supplemented with 1M ammonium sulfate at pH 7.2. Then we adjusted the ammonium sulfate concentration by mixing the buffer 1 and 2, accordingly. By applying different forces, we measured the step size or extension that described the unfolding and refolding of single protein domains. Gaussian fits were used to measure the absolute position and the required data corresponding to each step was obtained by length histograms. The differences in length between the centers of two peaks measured the step size.^3,6,7^

#### Image Processing

One of the most critical functions of magnetic tweezers is the determination of the Z position of the two beads, non-magnetic and paramagnetic. Non-magnetic beads were used as reference bead, stuck to the surface, to compensate the focal drift and instrumental vibration of the paramagnetic bead. To calculate the real time position of the bead, we used central processing unit (CPU) of the computer. We overclocked the CPU and enhance the image processing algorithm to get the satisfactory result.

There are several steps to determine the Z position of the bead:

1. Images are projected on a camera through objective that allows real-time screening of image connected to the computer and acquiring the region of interest (ROI).
2. Fourier transform (FFT) of the bead from the two ROI was acquired.
3. For determining the position of each bead, we calculate the radial profile and correlate to a Z-stack of the library and correlation profile was fitted.
4. The diffraction pattern around the bead can compute the change in end-to-end length of our protein of interest. We correlate measured radial vector and the stack at different position, for determining the z-position of the beads^3,8^.

#### Folding probability calculations

We represented the tables with residence time of each state at 10 pN force of figure 1 and calculated the folding probability using the eq. 1 mentioned in the result section.

**Supplementary Table 1:**
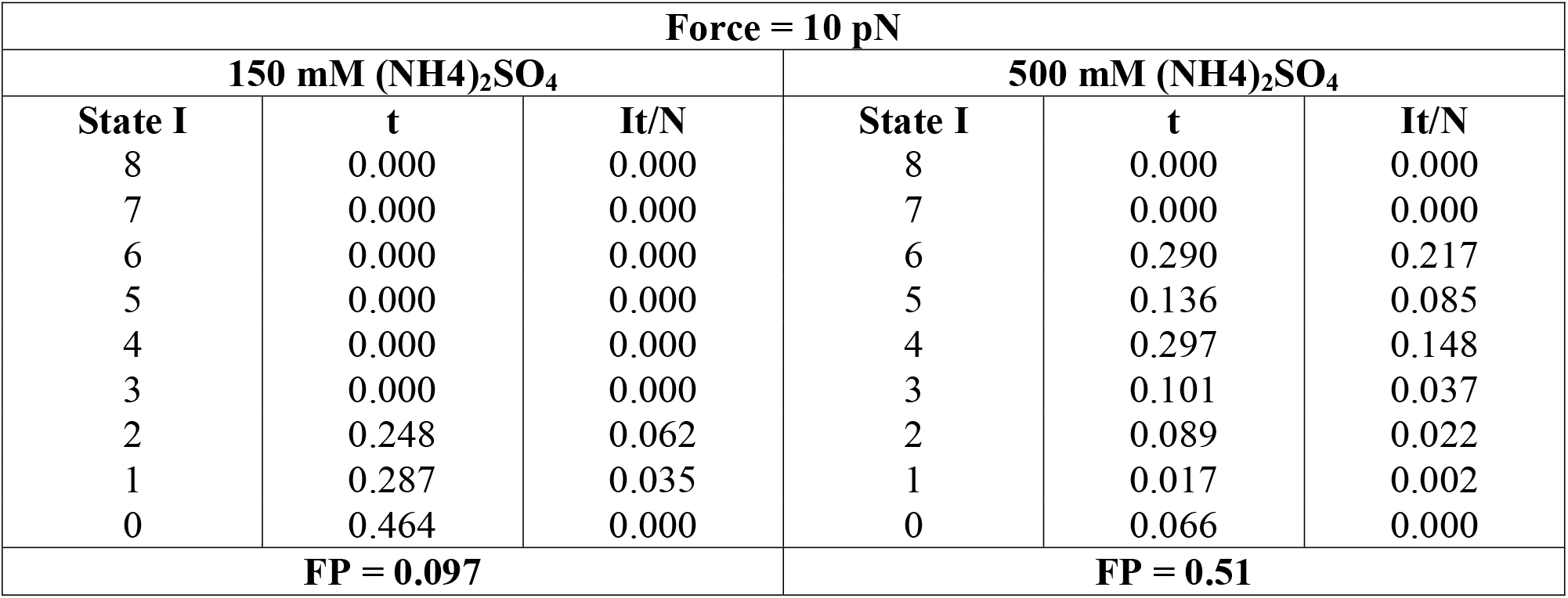
State occupation at 10 pN force with 150 and 500 mM (NH4)_2_SO_4_ of figure 1

#### Kuhn Length (*L*_*k*_) Calculations

We determined the Kuhn length for protein L by fitting the step size (SS) data in electrolyte freely jointed chain (Ec-FJC) model (eq. 6) against the varying forces at a particular salt condition.

**Supplementary Table 2:** Variation in Kuhn length with increasing salt concentration

| Salt Concentration (mM) | Kuhn Length, $L_k$ (nm) |
| --- | --- |
| 150 | 1.18±0.2 |
| 325 | 1.05±0.1 |
| 500 | 0.82±0.1 |
| 650 | 0.58±0.1 |

#### Protein L folding dynamics under varying salt conditions

We calculated the folding dynamics of protein L within 150 to 650 mM salt concentrations at 9 pN and observed salt shifts the folding dynamics of protein L towards the folded state and increases the FP from 0.41 at 150 mM to 0.97 with 650 mM concentration (Supplementary figure 2 to 5). These experiments were performed on different protein L molecules and are the supportive data to Fig. 2 (main text).

**Supplementary Figure 1:**
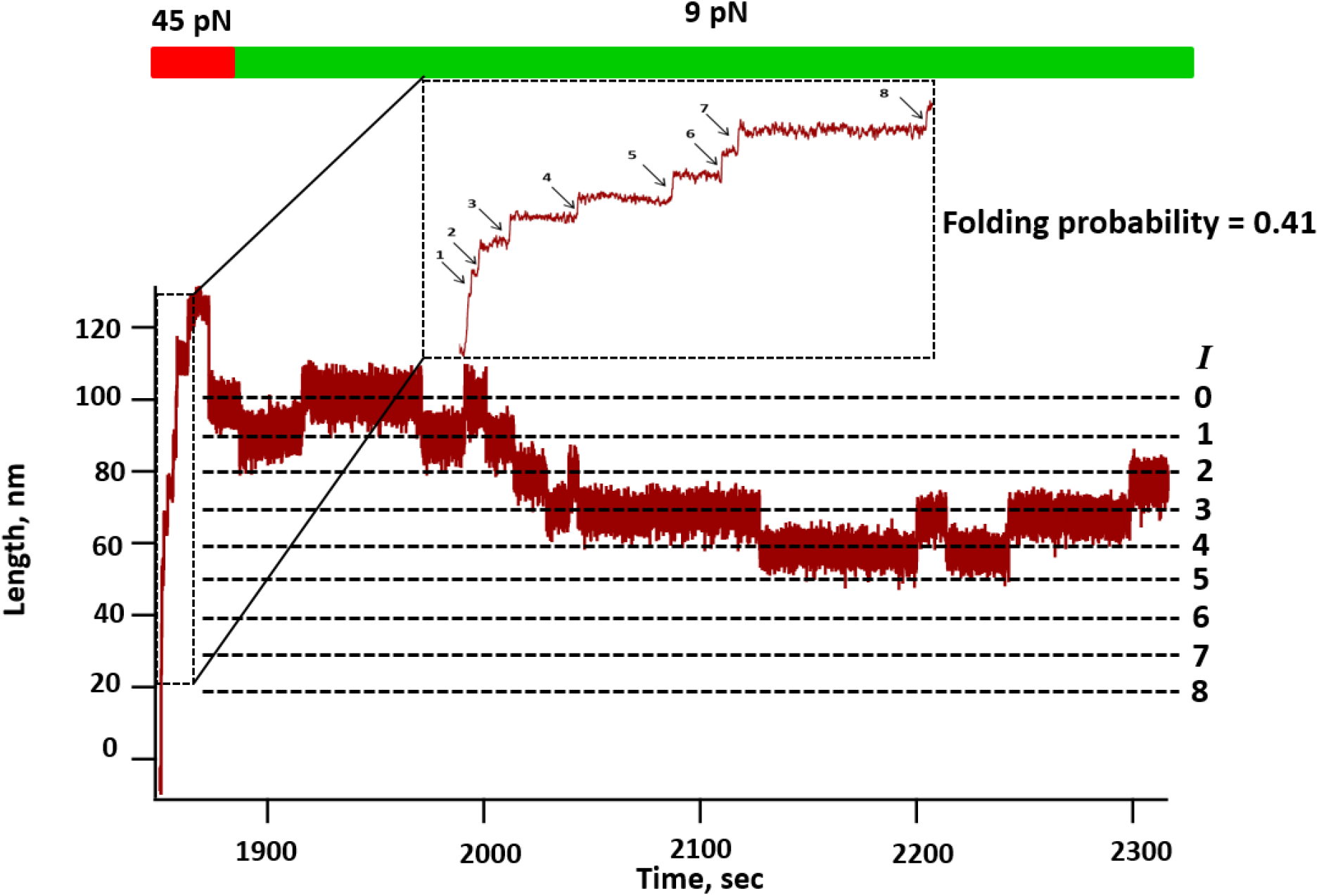
Effect of 150 mM (NH_4_)_2_SO_4_ on protein L folding dynamics: After full unfolding of protein L octamer at 45 pN, an equilibrium force of 9 pN was given in presence of 150 mM (NH_4_)_2_SO_4_ where the polyprotein exhibits dynamic folding-unfolding transition. Here the observed folding probability at 9 pN is 0.41.

**Supplementary Figure 2:**
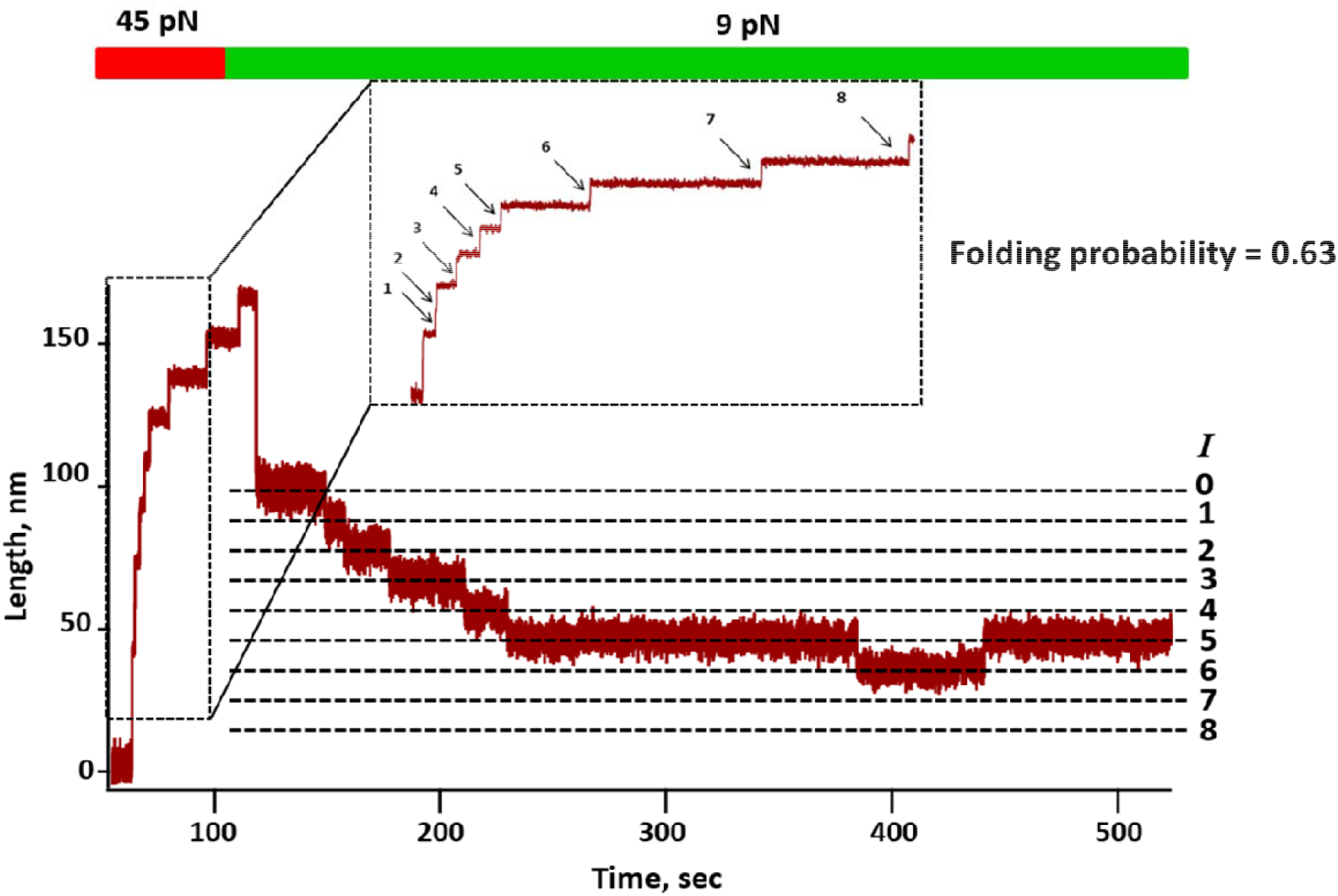
Effect of 325 mM (NH_4_)_2_SO_4_ on protein L folding dynamics: Unfolding force of 45 pN is applied on protein L, displaying eight discrete step sizes. Refolding pulse at 8 pN was provided to observe the folding dynamics at equilibrium, where we measured the folding probability is 0.63.

**Supplementary Figure 3:**
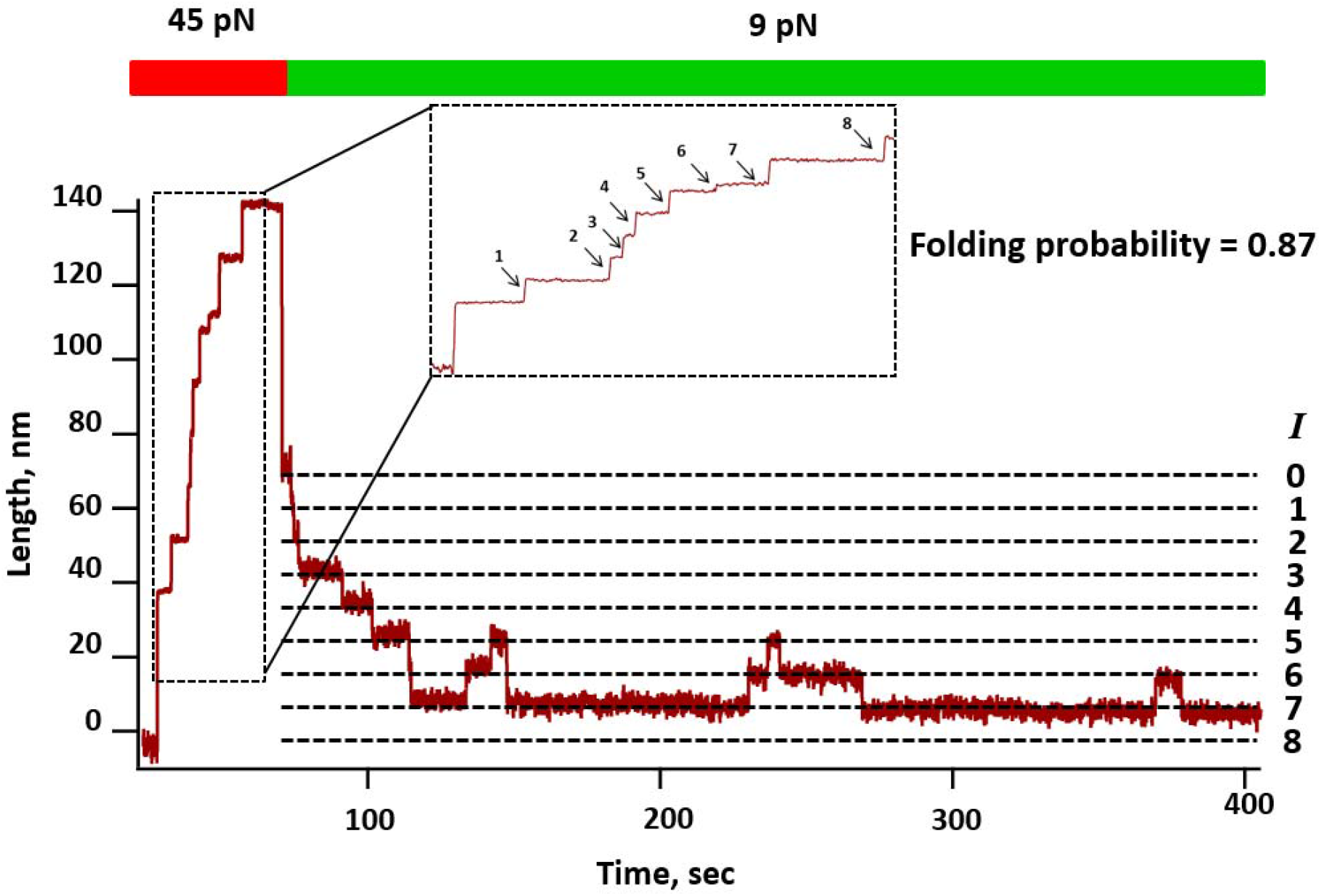
Effect of 500 mM (NH_4_)_2_SO_4_ on protein L folding dynamics: Initially, the polyprotein is fully unfolded at 45 pN force, getting eight unfolded domain that refolds back at 9 pN force where the folding probability has been observed to be 0.87.

**Supplementary Figure 4:**
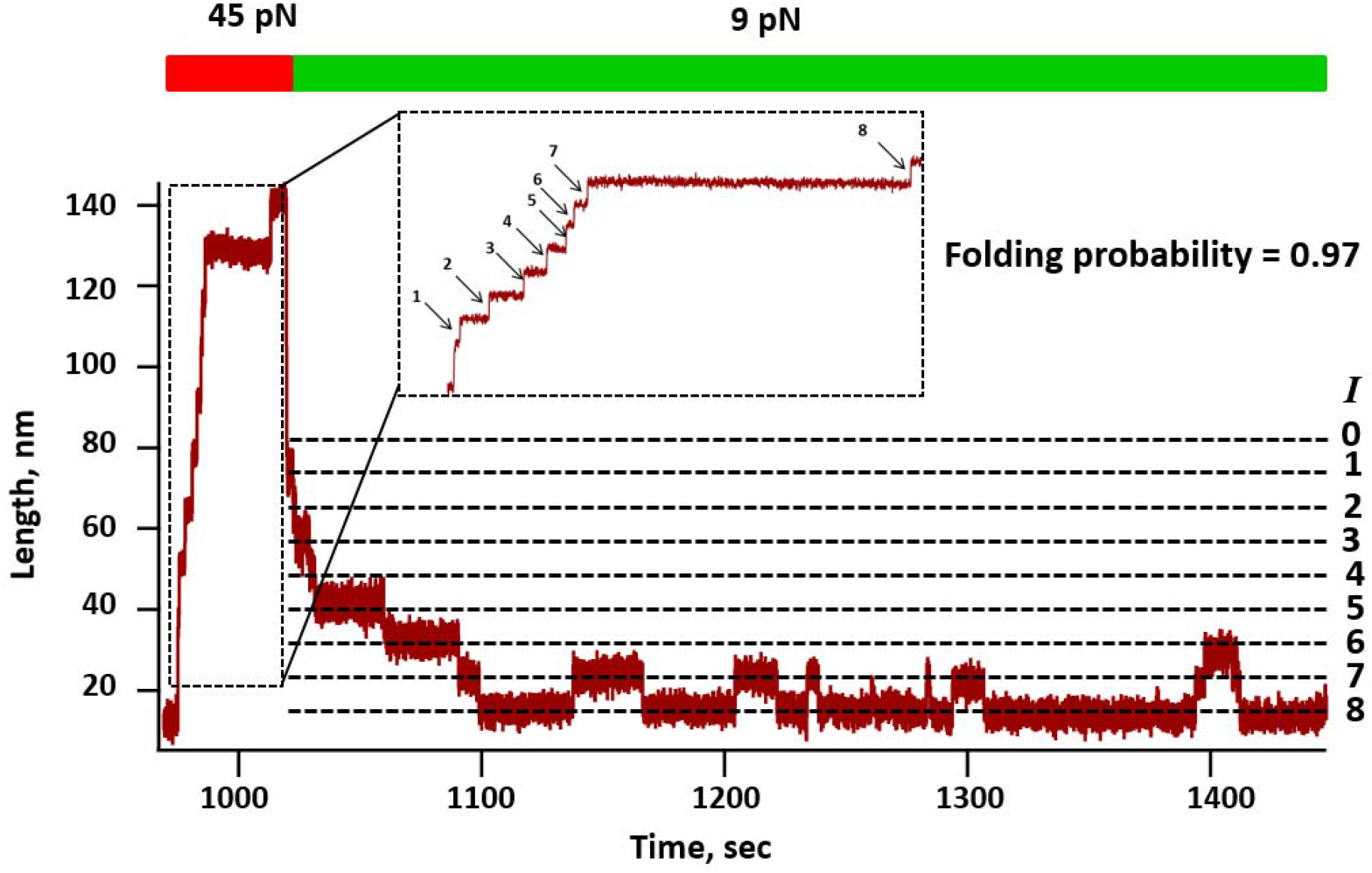
Effect of 650 mM (NH_4_)_2_SO_4_ on protein L folding dynamics: After applying an unfolding pulse at 45 pN, an equilibrium force of 9 pN was given in presence of 650 mM (NH_4_)_2_SO_4_ where the polyprotein exhibits dynamic folding-unfolding transition. The observed folding probability at 9 pN is 0.97.

**Supplementary Figure 5:**
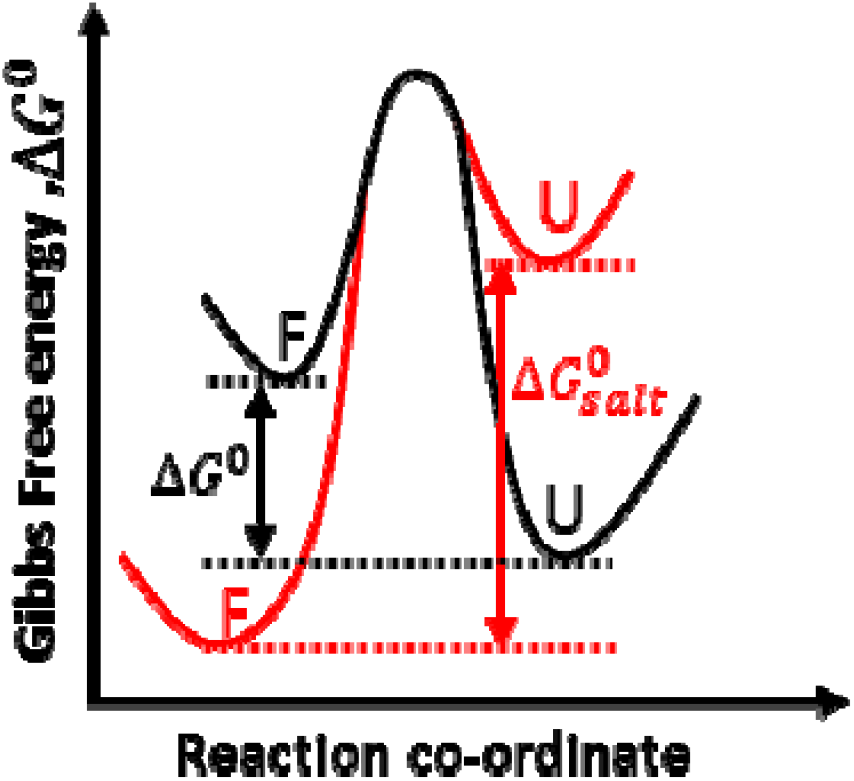
Salt reshapes the folding landscape of protein L: Salt changes the free energy landscape of protein L by stabilizing folded state and destabilizing unfolded state and thus, increases the difference in ΔG^0^ between folded and unfolded state. This leads to an increase in the unfolding free energy barrier; however, salt does not change the distance to the transition state.

**Supplementary Figure 6:**
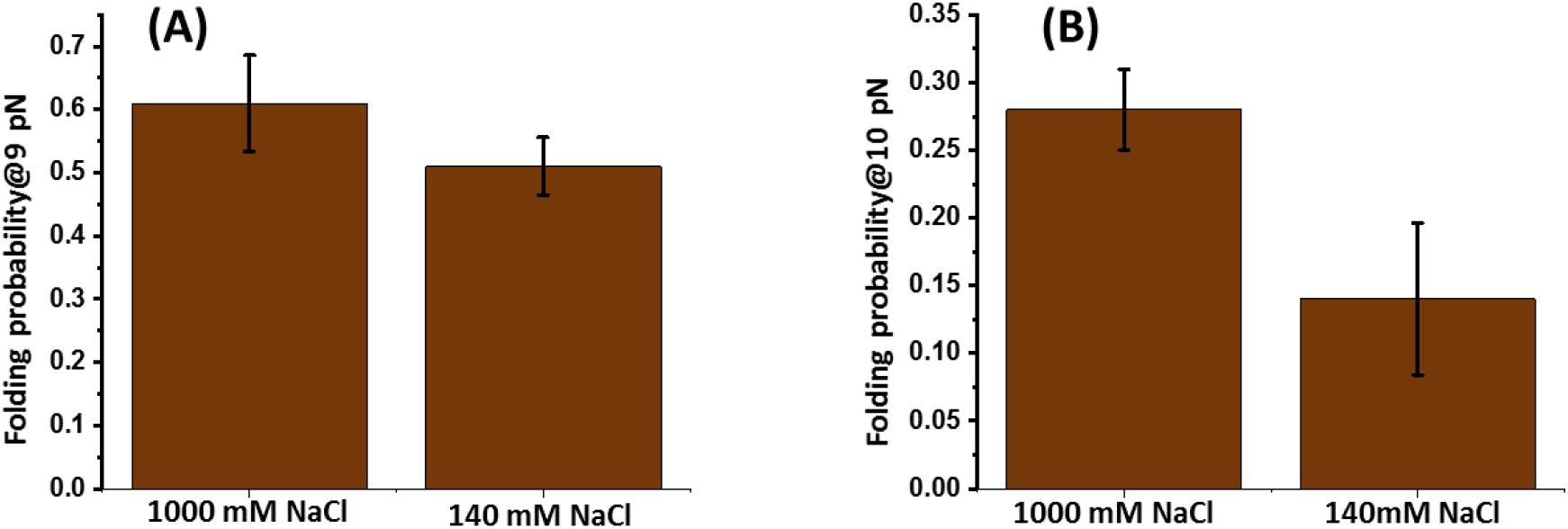
Variation of folding probability of protein L. **(A) at 9 pN:** With 140 mM salt, the folding probability is 0.61±0.07, while it decreases to only 0.51±0.05 in the presence of 1000 mM NaCl. **(B) at 10 pN:** Similarly, at 9 pN, with 140mM salt, the folding probability is 0.28±0.03 but reduces to only 0.14±0.06 at 1000 mM salt concentration. Both the Figure shows increase in folding probability with increase in salt concentration. Data points are calculated using >2500s and over six molecules per force. Error bars represent s.e.m.

**Supplementary Figure 7:**
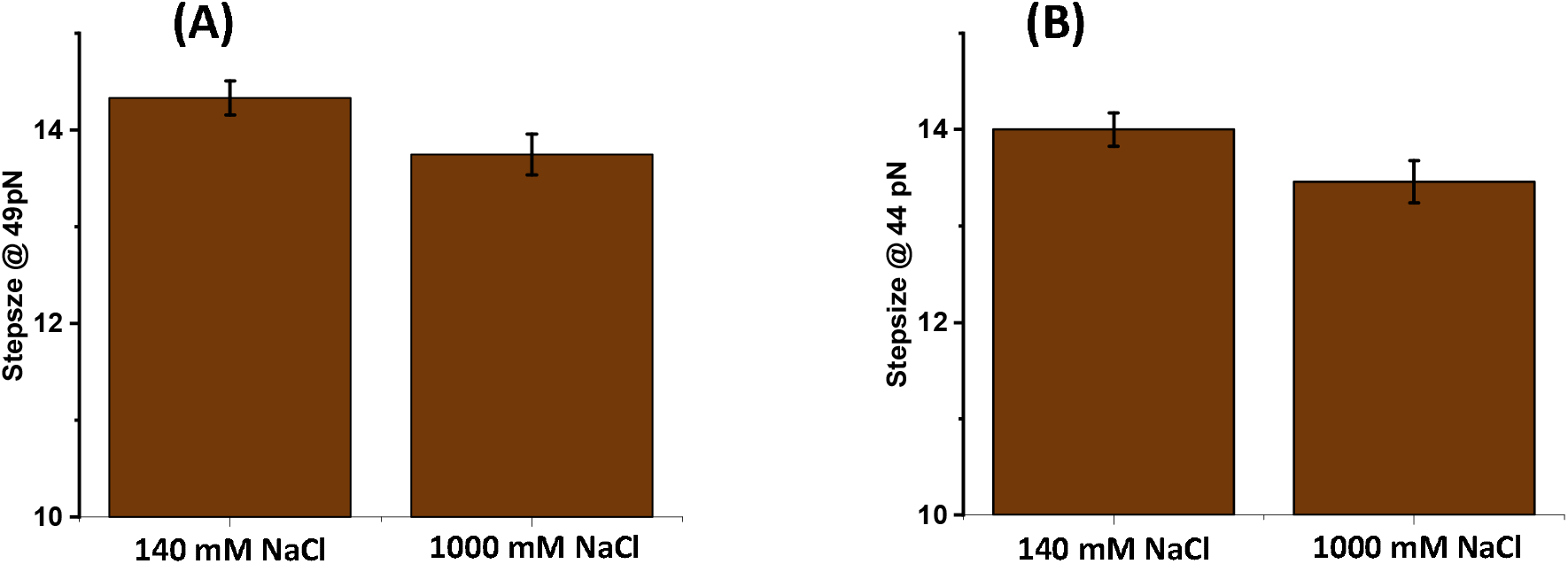
**(A) Step size difference of protein L at 49 pN:** In the presence of 140mM NaCl, the step size is 14.33±0.18 nm, while in the presence of 1000 mM NaCl, it decreases to 13.75±0.21 nm. **(B) Step size difference of protein L at 44 pN:** Similarly, at 44 pN, with 140 mM NaCl, the step size is14±0.18 nm which decreases to 13.46±0.22 nm at 1000 mM salt concentration. More than 10 molecules are measured and averaged for step size at different salt concentration. Error bars are s.e.m.

#### Force calibration of magnetic tweezers

In magnetic tweezers, the applied force can be empirically calibrated by several ways, depending on the construct length: either by measuring the lateral fluctuation for large construct (∼10 μm) such as nucleic acid or by relating protein unfolding extension to polymer elasticity models for smaller construct (∼50 nm).^9^ Since we are using small globular protein with step size with 6-15 nm, we sought to follow the calibration method for the protein construct.^6,7,10–16^

We have monitored the unfolding steps against the force in PBS buffer at 25ºC and fitted to the conventional freely jointed chain (FJC) model (Fig. 1) with the contour length of 16.6± 0.3 nm and Kuhn length of 1.1±0.2 nm. These values are in well agreement with the previous works by other groups^4,16^. To confirm the force calibration accuracy, we plotted the force at varying magnet distance and fitted to the exponential magnet law (Fig. 2).^3^

Furthermore, we have checked the sensitivity of force calibration by imposing the deviation values of protein L polyprotein construct: 0.3 nm for contour length and 0.2 for Kuhn length and observed that these length deviations affect the applied force within the 95% confidence level. Due to the bead-to-bead variation and difference in the tether attachment to the beads, the force has approximately 10% uncertainty.^3,17^ However, the M270 streptavidin beads from Invitrogen, have lower CoV of ∼2% than other dynabeads of Invitrogen, which could also negate the variation among dynabeads. However, for the proper force application, it is important to adjust the magnet distance by precisely regulating the linear voice coil.

#### Validating the force calibration accuracy by analyzing folding probability

We have validated our force calibration method by analyzing the folding probability (FP) data of protein L and then compared with protein L FP values, presented by different groups, who works on protein folding mechanics by magnetic tweezers.^6,7,11,18^ It is well-known that protein L exhibit strong force-dependent FP within 4 to 12 pN force, where the FP is ∼1 at 4 pN, while turns to zero at 12 pN. In our case, we observed that half-point force of protein L is ∼8 pN force and even 1 pN force deviation has strong effect on its half-point force. For example, at 7 pN, the protein L FP increases upto ∼56% (FP at 7 pN=0.8), while decreases by ∼72% at 9 pN (FP at 9 pN=0.14). Therefore, little alteration in the force calibration, due to the fluctuation of the magnet position and in the applied force, certainly perturb protein L FP under mechanical force. Interestingly, protein L exhibits half-point force at ∼8 pN force that are in well-agreement with their values (Fig. 3). Therefore, this force calibration method has a precise detection at sub-piconewton range, allowing us to observe the force-dependent folding dynamics, which strongly claims its fiduciary force measurement by our force calibration method. Lastly, the force calibration has also been confirmed by observing the conventional B-S overstretching transition of 565 bp dsDNA at ∼66 pN, which is within the range of well-defined force standard of DNA B-S transition at 65 pN.^19^ For the convenience of the reviewer, we have provided the figures below and also have included in the revised manuscript.

**Supplementary Figure 8:**
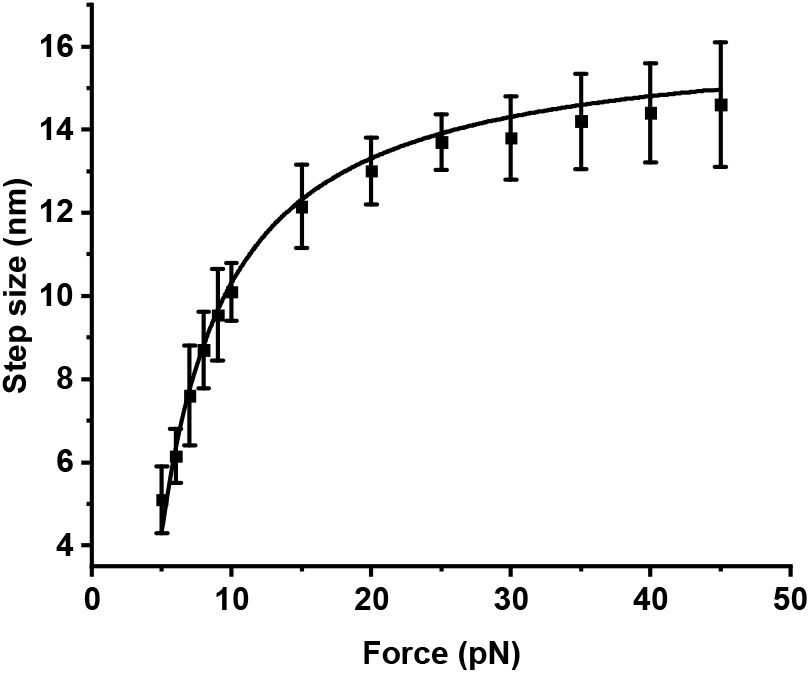
Step-sizes at different forces are fitted with freely jointed chain (FJC) model of polymer elasticity.

**Supplementary Figure 9:**
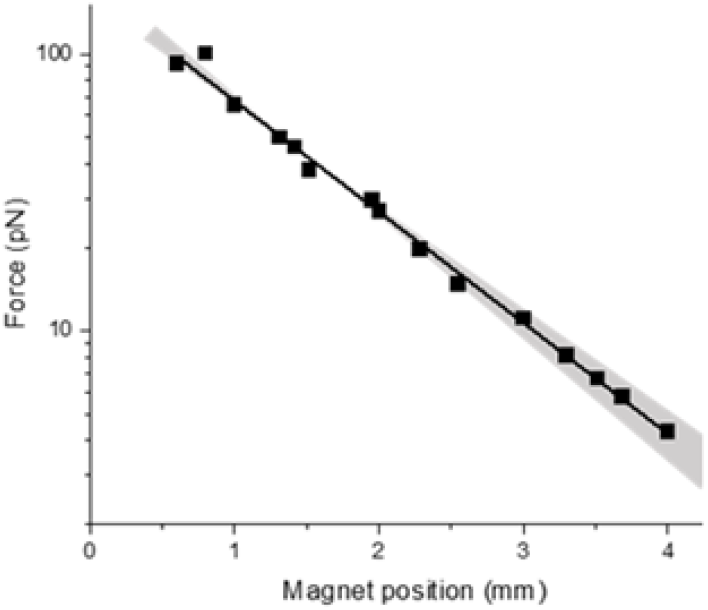
Force calibration by magnet law: We calibrated the force by the magnet law, which was proposed by Popa et al. 2016

**Supplementary Figure 10:**
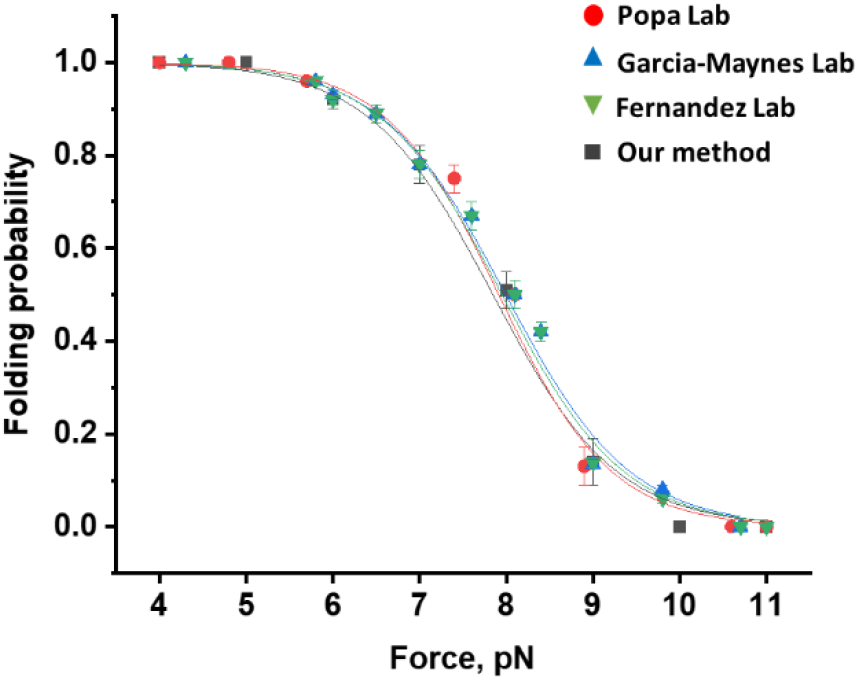
Confirmation of the force calibration by comparing folding probability of protein L: We have analyzed the folding probability by our force calibration method and compared with the folding probability values of protein L, studied by other groups.

